# A deep neural network provides an ultraprecise multi-tissue transcriptomic clock for the short-lived fish *Nothobranchius furzeri* and identifies predicitive genes translatable to human aging

**DOI:** 10.1101/2022.11.26.517610

**Authors:** Elisa Ferrari, Kathrin Reichwald, Philipp Koch, Marco Groth, Mario Baumgart, Alessandro Cellerino

**Author notes:** Contributing authors.

## Abstract

A key and unresolved question in aging research is how to quantify aging at the individual level that led to development of ”aging clocks”, machine learning algorhythms trained to predict individual age from high-dimensional molecular data under the the assumption that individual deviations of the predicted age from the chronological age contain information on the individual condition (often referred to as ”biological age”). A full validation of such clocks as biomarkers for clinical studies of ageing would require a comparison of their predictions with information on actual lifespan and long-term health. Such studies take decades in humans, but could be conducted in a much shorter time-frame in animal models. We developed a transcriptomic clock in the turquoise killifish *Nothobranchius furzeri*. This species is the shortest-lived vertebrate that can be cultured in captivity and is an emerging model organism for genetic and experimental studies on aging. We developed a proprietary deep learning architecture that autonomously selects a customizable number of input genes to use for its predictions in order to reduce overfitting and increase interpretability, and adopts an adversarial learning framework to identify tissue-independent transcriptional patterns. We called this architecture the Selective Adversarial Deep Neural Network (SA-DNN) and trained it on a multi-tissue transcriptomic dataset of *N. furzeri*. This SA-DNN predicted age of the test set with an accuracy of 1 day, i.e. less than 1% of the total species’ lifespan and detected genetic, pharmacological and environmental interventions that are known to influence lifespan in this species. Finally, a human transcriptomic multi-tissue clock that uses as input the orthologs of the genes selected by our SA-DNN in *N. furzeri* reaches an average error of ***∼*** 3 years rivalling epigenetic clocks. Our SA-DNN represents the prototype of a new class of aging clocks that provide biomarkers applicable to intervention studies in model organisms and humans.

---

Aging is the result of progressive and to some extent irreversible changes that impair cellular- and organ- physiology leading to an age-dependent increase in mortality risk. On a broad phylogenetic scale, interspecific differences in lifespan are the result of genetic differences that arose due to evolutionary processes. Interindividual variations in lifespan, on the other hand, are influenced by stochasticity [1] and environmental factors, such as nutrition [2] and only to a relatively minor extent by genetic variation, at least in humans [3]. A key and unresolved question in aging research is how to quantify aging at the individual level. Therefore, an increasing number of studies in the last decade focused on the development of machine learning methods, called aging clocks, trained to predict individual age from high-dimensional information, such as profiling of DNA methylation, gene expression, protein abundance etc. The great interest for such aging clocks arises from the assumption that individual deviations of the predicted age from the anagraphic age are not simple random errors, but contain also information on the individual condition. Subjects whose predicted age (often referred to as biological age) is higher than their calendar age have experienced age acceleration and are expected to be at higher risk of mortality and of developing age-associated illnesses. Age acceleration was indeed observed in response to pathological and physiological conditions [4–10] and -coversely-epigenetic age deceleration is currently regarded as a biomarker to assess the effects of interventions on human aging [11, 12]. However, other studies state that there is only a weak association between the biological age computed by these clocks and morbidity/mortality risk [13]. A full validation of such clocks as biomarkers for clinical studies of ageing would require to compare their predictions with information on the actual lifespan and long-term health. Such studies take decades in humans, but could be conducted in a much shorter time-frame in animal models, such as the mouse. In mice, verified life-prolonging manipulations do indeed have an impact on various DNA methylation based clocks [14–16], suggesting that these predictors can be used as biomarkers in interventional studies of aging. However, there is little overlap between the CpGs making up the mouse and human epigenetic clocks and the performance of the human CpGs on mouse age prediction is comparable to that of a random set of CpGs with the same numerosity[15], so they are not directly comparable. Therefore, the main objective of this work is to develop a molecular clock on an animal model, for which it is possible to easily test its sensitivity to environmental, genetic and treatment-based factors and to check its transferability to humans.

We decided to develop a transcriptomic clock because orthology greatly facilitates the transfer of clocks between species [17] and we focused on the turquoise killifish *Nothobranchius furzeri*. This species is the shortest-lived vertebrate that can be cultured in captivity and is an emerging model organism for genetic and experimental studies on aging [18–21], for which data are available on modulation of lifespan by genes, environment and pharmacological treatments. DNA methylation patterns proved to be particularly informative providing precise age prediction in a large set of human tissues [22, 23] and in cells from other mammals [24–28], transcriptomic clocks are less explored[17, 29, 30]. In particular, the success of the DNA methylation based clocks has been recently motivated by the discovery that cell- and tissue-independent age-associated DNA methylation changes are common [31]. Instead, the intrinsic tissue-specific nature gene expression, the differences in age-dependent transcriptome regulation across tissues [32], the dynamic nature of the transcriptome and the noise affecting its measurement may raise doubts on the possibility to develop a multi-tissue transcriptomic clock with performance comparable to its epigenetic counterparts. Therefore, our secondary objective is to explore the possibility to develop a multi-tissue transcriptomic clock with performance comparable to its epigenetic counterparts.

We choose to use a non-linear Deep Neural Network (DNN) approach to train our clock under the assumption that the most accurate transcriptomic pattern for age prediction would be specific for each tissue. DNNs gained popularity for their ability to extract features at different levels of abstraction and thereby learning more complex patterns [33]. Instead, most of the molecular clocks developed so far rely on the use of a linear model, the elastic net (EN), which is preferred over non-linear deep learning approaches, thanks to its resilience to overfitting and its interpretability. Besides the advantage that DNN can discover complex patterns and thus are more suitable for the development of a multi-tissue transcriptomic clock, some EN based studies have shown that introducing non-linear operations to the inputs [17] or to the outputs [34] of the model improves the prediction performance. This suggests that the aging process cannot be fully recapitulated by a linear combination of molecular data. However, DNN are notoriously affected by the so called confounding effects [35, 36], a phenomenon for which a model learns a predictive pattern based on a spurious association detected in the training set between the input data and the desired output. Therefore, our third objective was to design an algorithm that addresses overfitting, confounding effects and scarce interpretability while allowing to find a non-linear predictive pattern for accurate age prediction.

Thus, we developed our proprietary deep learning architecture that autonomously selects a customizable number of input data to use for its predictions in order to reduce overfitting and increase interpretability, and adopts an adversarial learning framework to avoid confounding effects. We called this architecture the Selective Adversarial Deep Neural Network (SA-DNN). We trained it on a multi-tissue transcriptomic dataset of *N. furzeri* and tested it on data collected under different lifespan-modulating conditions, being either due to genetic, environmental or treatment factors. Finally, we assessed whether the human orthologs of the genes selected by our SA-DNN for the age prediction in *N. furzeri* are informative also for a human transcriptomic multi-tissue clock. Our analyses allow us to draw general conclusions on the usefulness of these clocks to predict biological age.

## 1 Results

### 1.1 Study design

For this study we analyzed five RNA-seq datasets of *N. furzeri* :

1. A multi-tissue dataset comprising transcriptomic profiles of brain, liver, skin and muscle samples of 179 fishes, belonging to 3 different strains that have a life expectancy of ∼ 8-9 months in our facility (see Fig. 11) for a total of 308 samples, 100 sequenced in the present study.
2. A public longitudinal dataset comprising transcriptomic profiles of fin biopsies collected from 150 fish at 10 and 20 weeks of age. All the fish analyzed in this dataset belong to the MZM0410 strain, that showed a median lifespan of 39 weeks [37].
3. A dataset comprising transcriptomic profiles of brain, liver and skin samples of fish belonging to the GRZ strain, which has a life expectancy much shorter than the strains composing the previous two datasets *∼* 20 weeks in our facility (see Fig. 11), for a total of 147 samples, 72 sequenced in the present study.
4. A public transcriptomic dataset comprising fish treated with rotenone for four weeks before being sampled at age 9 or 27 weeks and the respective untreated controls. A previous study on this dataset has shown that rotenone treatment at these two ages elicits opposite effects on lifespan [38].
5. A public dataset containing 26 transcriptomic brain profiles of *N. furzeri* collected in the wild where they display faster growth and shorter life expectancy with respect to captive ones[39, 40].

The most important information on these datasets, according to the analyses conducted, is summarized in the Tables 1, 2, 3, 4, 5. Details about dissection, gene expression profiling and RNA-seq quantification can be found in Sec. 4.1 and 4.2.

**Table 1:** Dataset 1 phenotypic composition.

| **(1) Multi-tissue dataset**<br>**Multi-tissue dataset of long-living strains** |  |  |
| --- | --- | --- |
| Sample | 308 |  |
| Fish | 179 |  |
| Tissue | Brain | 129 |
|  | Liver | 55 |
|  | Skin | 109 |
|  | Muscle | 15 |
| Strain | MZM0410 | 192 |
|  | MZCS08122 | 52 |
|  | MZM0403 | 64 |
| Sex | Male | 276 |
|  | Female | 32 |
| Housing | Single | 253 |
|  | Group | 55 |
| Batch | 12 different batches |  |

**Table 2:** Dataset 2 phenotypic composition.

| **(2) Fin logitudinal dataset**<br>**Longitudinal finclip dataset of a long-living strain** |  |  |
| --- | --- | --- |
| Samples | 300 |  |
| Fishes | 150 |  |
| Sample age | 10 weeks | 150 |
|  | 20 weeks | 150 |
| Tissue | Fin |  |
| Strain | MZM0410 |  |
| Sex | Male |  |
| Batch | batch1 | 90 |
|  | batch2 | 210 |
| Hatch date | 11 dates between 2012 and 2014 |  |

**Table 3:** Dataset 3 phenotypic composition.

| **(3) GRZ dataset**<br>**Multi-tissue dataset of short-living strains** |  |  |
| --- | --- | --- |
| Samples | 147 |  |
| Tissue | Brain | 61 |
|  | Liver | 61 |
|  | Skin | 25 |
| Strain | GRZ |  |

**Table 4:** Dataset 4 phenotypic composition.

| **(4) Rotenone's dataset** |  |  |
| --- | --- | --- |
| Samples | 69 |  |
| Sample age | 9 weeks | 24 |
|  | 27 weeks | 45 |
| Tissue | brain | 25 |
|  | liver | 22 |
|  | skin | 22 |
| Strain | MZM0410 |  |

**Table 5:** Dataset 5 phenotypic composition.

| (5) Wild fishes dataset |  |
| --- | --- |
| Samples | 26 |
| Tissue | Brain |
| Batch | 2 different batches |

The multi-tissue dataset (dataset 1) was used to train, validate and test the Nothobranchius’ Transcriptomic Intelligent ClocK (N-TICK). The longitudinal dataset (dataset 2) was used as an independent test to verify the generalizability of the N-TICK on a tissue not used for training (i.e. the fin). The remaining three sets were used as independent test sets to verify if NTICK predictions are influenced in the expected direction by conditions known to influence lifespan of *N. furzeri*. To study genetic effects, we applied it to the short-lived GRZ strain (dataset 3). To study pharmacological modulation of lifespan we used the Rotenone’s dataset (dataset 4) and to test environmental effects we used the wild dataset (dataset 5).

In addition, a reduced version of the N-TICK, called N-rTICK, has been developed by making our SA-DNN more selective, meaning that it is forced to select a reduced number of input data to use for inference. The N-rTICK was trained on the multi-tissue dataset (dataset 1) and tested on all the other datasets, as already described for the N-TICK.

Finally, to test if the genes selected by our N-TICK and N-rTICK are informative also of the aging process of humans, we trained a Human Transcriptomic Intelligent ClocK (H-TICK) using as input only the human orthologs of these genes. For this operation, we used the public database of Genotype-Tissue Expression (GTEx) project, [41]. GTEx comprises the transcriptomic profiles of 54 different non-diseased tissues taken post-mortem from 948 donors with an age span of 20-71 years. Given that the information available on the age of the subjects was reported as age classes spanning 10 years, we also trained a modified version of the H-TICK, called H-cTICK, specifically designed to address the prediction as a classification problem instead of the typical regression approach applied to generate aging clocks. The most important information on the GTEx dataset, according to the analyses conducted, is summarized in Table 6.

**Table 6:** GTEx dataset phenotypic composition.

| (6) Human GTEx dataset |  |  |
| --- | --- | --- |
| Samples | 17225 |  |
| Subjects | 932 |  |
| Tissues | 27 |  |
| Nucleic acid isolation batches | 1839 |  |
| Nucleic acid isolation batches | 565 |  |
| Sex | Male | 11506 |
|  | Female | 5719 |
| Type of death | 0 - during ventilation | 8777 |
|  | 1 - fast, violent | 710 |
|  | 2 - fast, natural | 4835 |
|  | 3 - after terminal phase of 1-24 hours | 867 |
|  | 4 - slow after long illness | 2036 |

### 1.2 The Transcriptomic Intelligent ClocKs (TICKs)

We developed a new algorithm, called Selective Adversarial Deep Neural Network (SA-DNN), to address the overfitting and the confounding problems while searching for a non-linear but to some extent interpretable pattern. In summary, the SA-DNN core features are the use of an initial selector layer and the adoption of an adversarial learning framework.

The initial selector layer is a one-to-one binary layer assigning the weight 0 or 1 to each input feature, to select only those most informative for the aging prediction and thereby mitigating the overfitting problem. The selector layer is regularized in order to make it possible for the user to influence the number of features to select by changing the value of one hyperparameter (hereinafter referred to as *”t”*). The input features that were selected and used to make the predictions are those showing weight =1 in the selector layer after the training session. So, the selector layer reduces the overfitting and increases the interpretability of the model by greatly reducing the number of input genes.

The adversarial learning framework, instead, is used in our SA-DNN to minimize the confounding effect of a set of variables, defined by the user. The SA-DNN is then trained to converge on the best trade-off between minimizing the error on the desired task, i.e. the age prediction, and maximizing the error on the identification of the variables selected by the user that may mislead the learning process. This process ensures that the SA-DNN has learned to predict the task of interest specifically and it has not learned a pattern based on spurious correlations between the outcome and some confounding variables. A more detailed description of the SA-DNN is provided in Sec. 4.4.

In this work the SA-DNN was used to develop 4 different clocks, called N-TICK, N-rTICK, H-TICK and H-cTICK. The first two are trained on the same set of *N. furzeri* data from the multi-tissue dataset (dataset 1) to predict the age of the fish at sampling. Both receive as input the expression of 19,711 genes expressed in all the different tissues composing the dataset (i.e. the genes with more than 10 counts in at least 80% of the samples for all tissues). The variables set to be countered by our adversarial learning framework are: tissue, sex, strain, housing (i.e. whether the animal was raised in individual- or community-tank), sequencing batch and the identifier of the individual fish, since different tissues were sampled from the same individual. The N-rTICK differs from the N-TICK only for the number of selected genes. In particular, the former was developed as a reduced version of the N-TICK to test the effects on accuracy of a reduction in the number of genes used to train the age predictor.

The H-TICK and the H-cTICK are trained on a set of human data from the GTEx database (dataset 6) to predict the age of the samples. However, given that the dataset provides the age labels only using 10-year spanning classes, we applied two different approaches. The H-TICK was developed as a regressor trained to predict the middle value of each age class reported, while the H-cTICK was trained as a classifier. With this second approach, the classifier is equally penalized when it erroneously predicts any class other than the real one during the training session, irrespectively of the age difference between the two classes. Methodological difference aside, both the H-TICK and H-cTICK were developed to test if the genes selected by the N-TICK and N-rTICK were also informative of the human aging process. Thus, the two clocks have been designed to receive as input the expression of the human orthologs only of the *N. furzeri* genes selected by N-TICK and N-rTICK and their selector layer has been inactivated. The GTEx is a highly annotated dataset and the information of various possible confounding variables are available. However, due to the wide number of samples available we decided to correct the effect of only the few variables that, in our opinion, may affect the gene expression values the most, which are: tissue, sex, type of death and subject identifier. A summary of the 4 clocks is provided in Tab. 7.

**Table 7:** Summary of the developed clocks.

| Clock | Dataset | Input genes | Selected genes | Confounders | Output |
| --- | --- | --- | --- | --- | --- |
| N-TICK | (1) | 19,711 | 420 | Tissue<br>Strain<br>Sex<br>Housing<br>Batch<br>Fish ID | Age<br>(continuous range) |
| N-rTICK | (1) | 19,711 | 80 | Tissue<br>Strain<br>Sex<br>Housing<br>Batch<br>Fish ID | Age<br>(continuous range) |
| H-TICK | (6) | 293 | All* | Tissue<br>Sex<br>Type of Death<br>Subject ID | Age<br>(continuous range) |
| H-TICK | (6) | 293 | All* | Tissue<br>Sex<br>Type of Death<br>Subject ID | Age<br>(10 years intervals) |

**Table 8:**
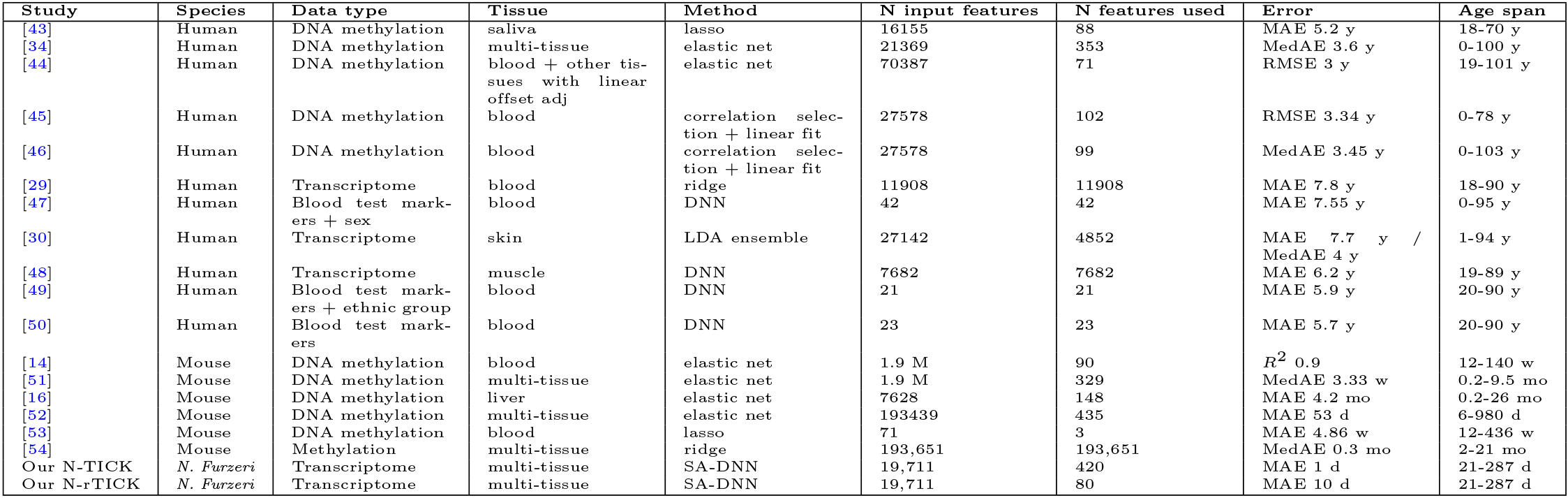
Summary of the results obtained by other clocks in literature. The 6th column shows the dimensionality of the input data (i.e. the number of features analyzed) and the 7th one shows the number of input features actually used in the inference process. The last two lines reports the results obtained in this study with ours N-TICK and N-rTICK.

An exhaustive description of the methodological choices made to design, train, validate and test the clocks can be found in Secs. 4.4 and 4.5. To avoid overestimation of the clock’s performance, the training, validation and test sets always contained data belonging to different individuals [42] for both the human and fish datasets.

### 1.3 N-TICK provides an ultraprecise multitissue age prediction for *N. furzeri*

We tested if our SA-DNN architecture is able to discover predictive tissue-independent age patterns in transciptome data by training the N-TICK predictor, setting the tissue identifier as one of the confounding variables, in order to make the learning process independent from this variable. Fig. 1 shows the results obtained by applying the SA-DNN on the datasets 1 and 2. The Root Mean Squared Error (RMSE) and the Mean Absolute Error (MAE) are *∼* 1-2 days for the training, validation and test sets obtained partitioning the multi-tissue dataset (dataset 1) 1. The magnitude of the prediciton errors does not increase when the N-TICK is applied to the fin longitudinal dataset (dataset 2), which is composed solely by a tissue not represented in dataset 1 (hold-out set). This last observation supports the notion that our adversarial approach enables the development of a generalizable and tissue-independent clock. As a confirmation, we report in Appendix A.3 an analysis of the distribution of the residuals across various categories stratifying the population of datasets 1 and 2, including the tissue type. This analysis confirms the small size of the errors and reveals their even distribution across the different categories. In addition, Appendix A.2 shows that in the absence of our adversarial learning framework, the hidden features identified by a DNN and used for age prediction are dependent on all the variables that were penalized by the SA-DNN, with tissue type having the largest effect.

**Fig. 1:**
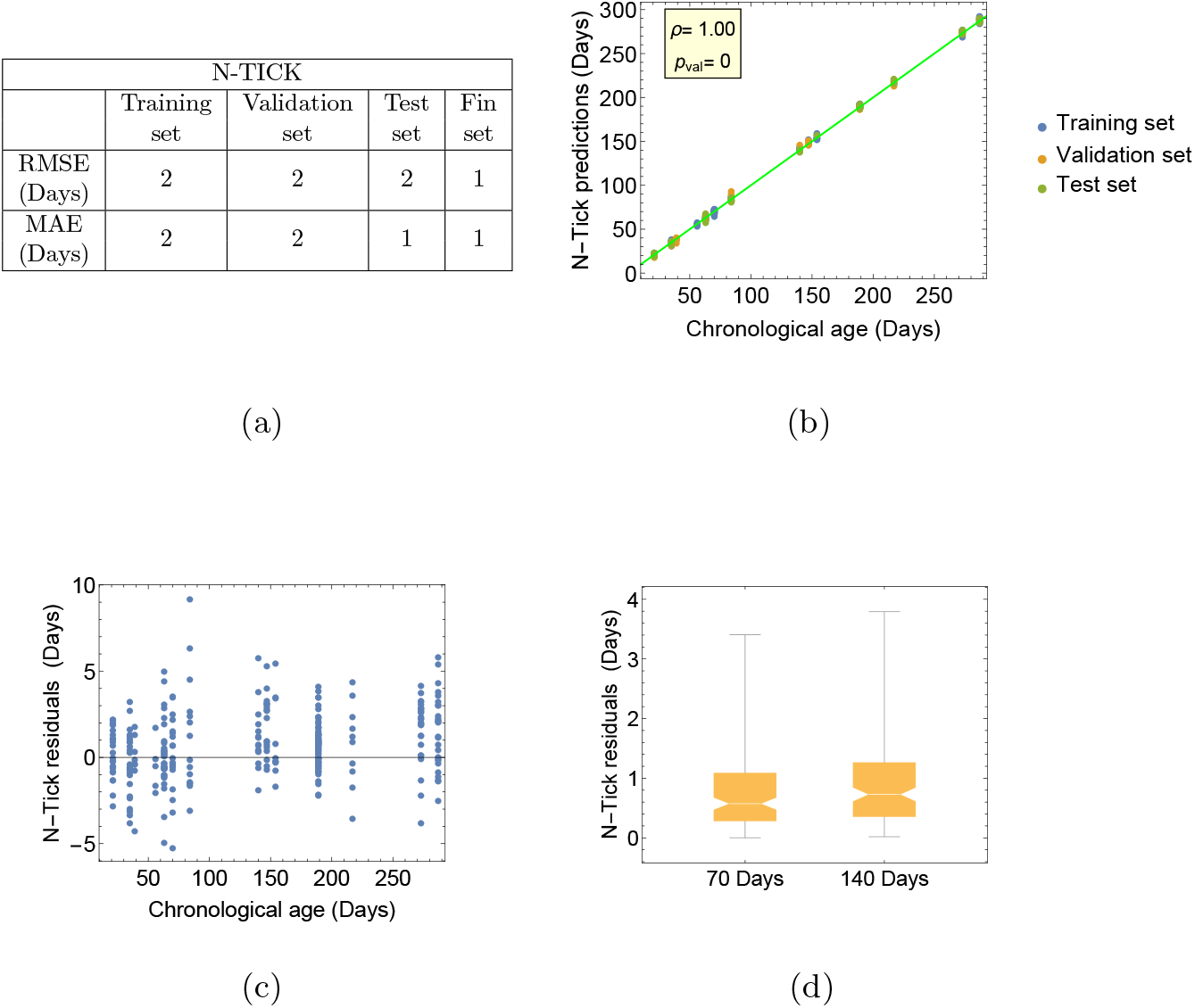
Performance of the N-TICK on the multi-tissue and longitudinal datasets (i.e. the datasets 1 and 2). Fig. (a) reports the RMSE and the MAE computed on the training, validation and test sets defined from the multi-tissue dataset, described in Table 1. The last column shows the performance obtained on the hold-out longitudinal dataset composed by fin samples, described in Table 2. Fig. (b) shows the correlation between the predicted and the true chronological age when applying N-TICK to the multi-tissue dataset. Fig.s (c) and (d) show the residuals of the N-TICK predictions with respect to chronological age on the multi-tissue and longitudinal datasets, respectively.

In addition to the excellent performance obtained, the errors are evenly distributed with respect to age Figs. 1b-d and the performance remains stable across the training, validation, test and hold-out sets (Fig. 1), indicating that N-TICK did not overfit the data. As already mentioned in Sec. 1.2, the overfitting problem was addressed by our SA-DNN by the introduction of the selector layer, whose hyperparameter *t* was set to select a number of genes in the same order of magnitude of the features used by other multi-tissue clocks in literature (around 320-440 features [34, 51, 52]). The number of genes selected by our N-TICK is 420, which eases potential performance comparisons with other methods for age prediction described in the literature.

Summarizing, we developed a multi-tissue clock based on the expression of 420 genes using our SA-DNN architecture. This resulting clock is unaffected by overfitting and its errors are independent from age as well as close to their theoretical lower limit.

### 1.4 N-TICK is sensitive to genetically-determined differences in aging rates

To test if N-TICK is sensitive to genetically-determined differences in aging rates, we applied it to dataset 3. This dataset comprises solely samples of individuals of the GRZ strain whose lifespan in captivity is roughly one half than that of the strains used to train the N-TICK. The age of the GRZ samples are significantly overestimated and are fitted by a line with a slope almost equal to 3 (see Fig. 2b), implying that the GRZ strain shows a transcriptomic age acceleration of a factor 3 according to N-TICK predictions. The errors with respect to this fit (Fig. 2a-b) are significantly larger than those observed on datasets 1 and 2; they are non-randomly distributed and increase with age. This suggests that the age-dependent patterns of transcriptome regulation in this shorter-living strains of *N. furzeri* are not simply scaled, but also are influenced to some extent by idiosyncratic genetic effects. However, a 95% correlation between true age and predicted age (defined as the best linear fit of the data) remains remarkable, considering that the samples belong to a different strain with respect to those used to train the model.

**Fig. 2:**
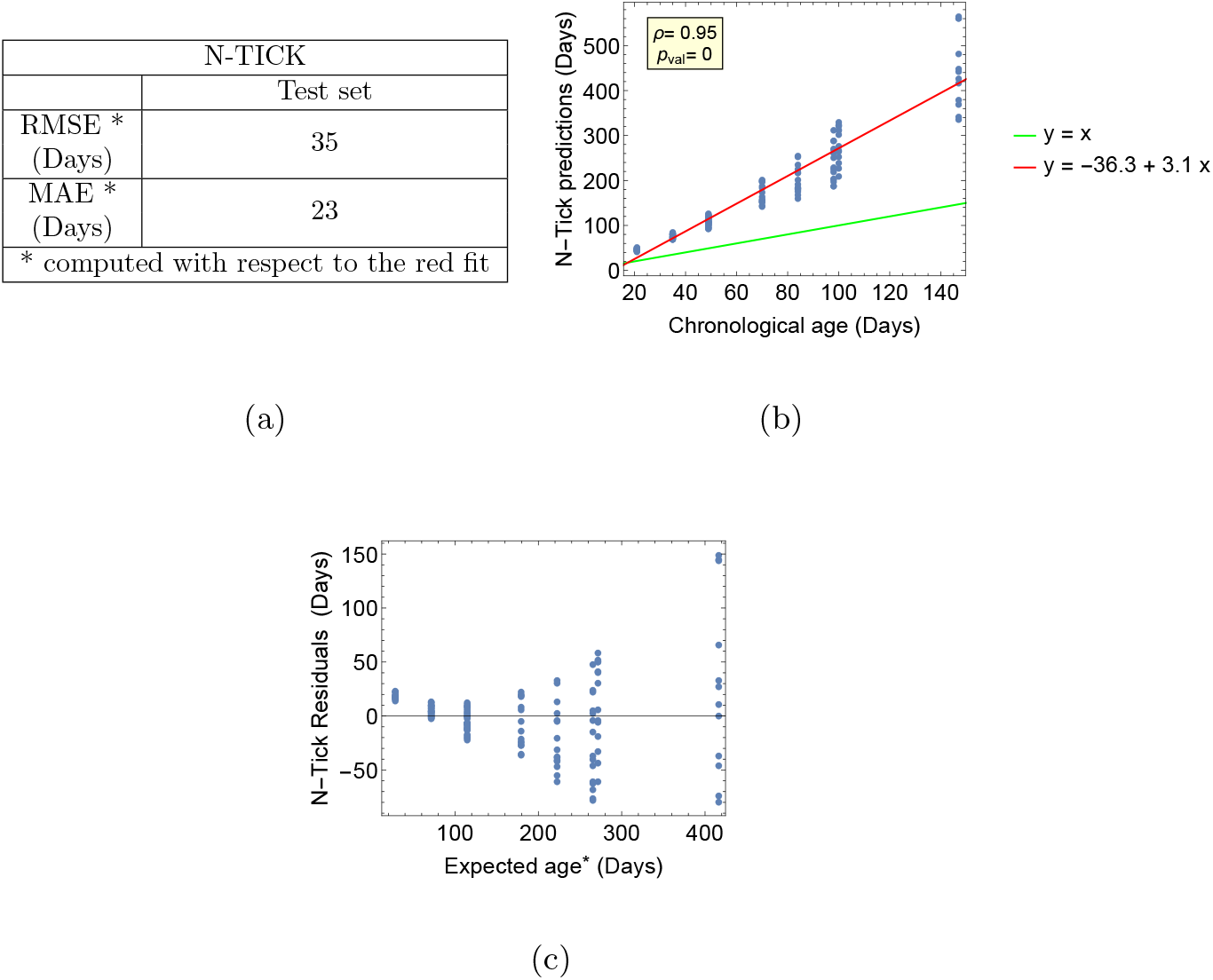
Performance of the N-TICK on the GRZ datasets (i.e. the dataset 3). Fig. (a) reports the the RMSE and the MAE tested on the GRZ dataset. Note that to capture the spread of the errors instead of their shift due to the aging rate difference between the strains used in the training set and the GRZ one, the RMSE and the MAE are computed with respect to the expected age fit by the red line in Fig. 2b. Fig. (b) shows the correlation between the predicted and the true chronological age when applying N-TICK to the GRZ dataset. Fig. (c) show the residuals of the N-TICK predictions with respect to the expected age (i.e. the one fit by the red line in Fig. 2b) on the GRZ dataset.

### 1.5 N-TICK is sensitive to environmentally-determined differences in aging rates

To test if the N-TICK developed is also sensitive to environmentally-determined differences in aging rates, we compared its predictions on wild and captive fishes, using the data of the wild fishes dataset (dataset 5), which comprises solely brain samples, and the brain samples of the multi-tissue dataset (dataset 1). The growth rate and maturation of *N. furzeri* is faster in wild habitats than in captivity when population density is low, which is the case for the population that constitutes dataset 5. [39, 40].

Accordingly, the predicted age for the wild brain samples is consistently higher than their true age Fig.s 3a-b, while the predictions of the captive brain samples are very accurate. The spread of the errors on the wild samples is larger than that of the captive ones. This reduction in precision is expected since wild animals show basal differential expression of almost one third of expressed genes as compared to captive-raised fish and some genes are regulated in opposite directions by age in the two conditions [55]. Furthermore, it should be considered that a constant source of error in the age of the wild fish arises from the uncertainty in their age estimation that is obtained from daily otolith increments [56]. It can be also noted that the spread of the errors on the wild samples increases with age, this phenomenon could be explained by the accumulation of environmental effects on gene expression that causes divergence of individual phenotypic trajectories. Wild samples of *N. furzeri* of known age are difficult to obtain and dataset 5 allows us to explore the effect of the N-TICK only on a limited age range with only three points. Within this age range, the wild fish show higher transcritptomic age but lower age acceleration with respect to conspecifics raised in captivity. However, the aging rate of the wild fish could be non-linear with respect to the pattern learned on the captive ones.

**Fig. 3:**
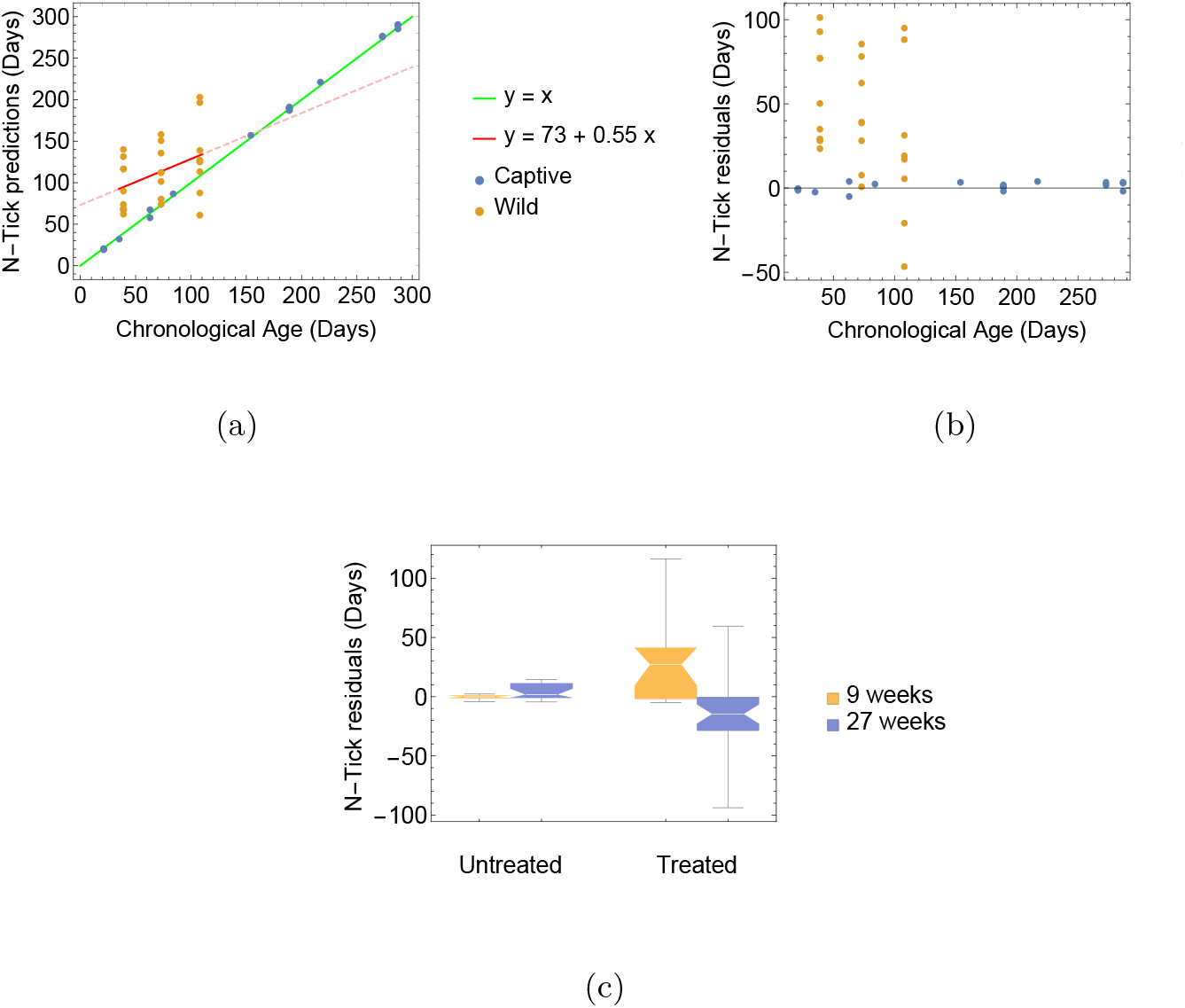
Comparison of the N-TICK predictions obtained on wild and captive animals (Fig.s a-b) and on individuals untreated and treated with Rotenone (Fig. c). Fig. (a) shows the N-TICK predictions with respect to the true chronological age of wild and captive brain samples of the MZM strain. The brain samples of the captive fishes (blue dots) are taken from dataset 1, while the ones of the wild individuals (orange dots) are taken from dataset 5. Fig. (b) shows the residuals of the predictions illustrated in Fig. a. Fig. (c) shows the residuals of the N-TICK predictions on samples untreated and treated with Rotenone at two different ages. The data for this experiment are taken from dataset 4.

### 1.6 N-TICK detects anti-aging and pro-aging effects of a small compound

To test the ability of N-TICK to capture drug-induced modulations of lifespan, we applied it to the Rotenone’s dataset (dataset 4), which comprises the data of fish treated for 4 weeks with 15 pM of Rotenone in the water and sampled at two different ages (9 and 27 weeks) and their respective controls. A previous study on *N. furzeri* observed that treatment with this concentration of Rotenone induces opposite effects on lifespan at these ages, causing life-extension in older fish and life-shortening in young fish [38]. As Fig. 3c shows, the life-span modulating effects of Rotenone observed with an *in vivo* experiment are also detected in-silico by our clock, which overestimates and underestimates the age of the treated fishes at 9 and 27 weeks, respectively, consistently with the experimentally-measured effects on lifespan. The ages of the control groups, instead, are predicted with a very small error in the range of the errors reported in Fig. 1 for the multi-tissue clock, as a further confirmation of the accuracy of N-TICK.

### 1.7 Effects of reducing predictor gene numbers on accuracy

We explored different configurations of our SA-DNN in order to identify the minimum number of input genes necessary to obtain an age prediction error not exceeding 2 weeks by modifying the hyperparameter *t* that influences the number of features selected by the filtering layer. Following this strategy, we developed a reduced version of the N-TICK, called N-rTICK, that uses only 80 genes as input. Its performance on the muti-tissue and longitudinal datasets is shown in Fig. 4. As expected, reducing the number of input genes reduces the accuracy, yet the correlation between the predictedan the true-age is still *ρ* = 0.99 (Fig. 4a). Notably, in this reduced version of the clock, the errors are not random and increase with age (Fig.s 4c-d). Also in this case, we do not detect signs of overfitting, i.e. larger errors in the test- and validation-set as compared with the training-set (Fig. 4a).

**Fig. 4:**
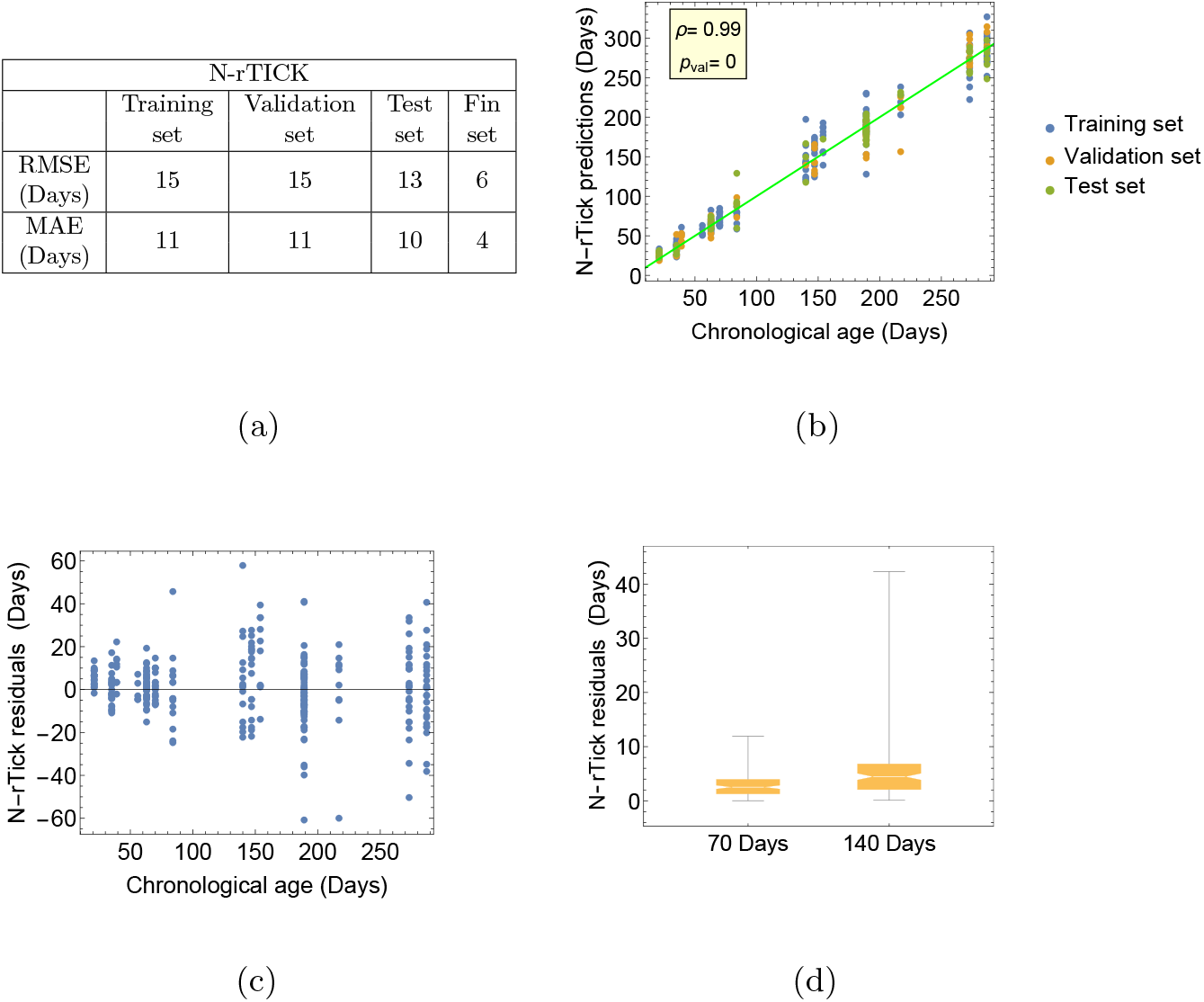
Performance of the N-rTICK on the multi-tissue and longitudinal datasets (i.e. the datasets 1 and 2). Fig. (a) reports the the RMSE and the MAE computed on the training, validation and test sets defined from the multi-tissue dataset, described in Table 1. The last column shows the performance obtained on the hold-out longitudinal dataset composed by fin samples, described in Table 2. Fig. (b) shows the correlation between the predicted and the true chronological age when applying N-rTICK to the multi-tissue dataset. Fig.s (c) and (d) show the residuals of the N-rTICK predictions with respect to chronological age on the multi-tissue and longitudinal datasets, respectively.

In Appendix A.4, we repeated the analyses described in Sec.s 1.4-1.6 using the N-rTICK to test whether this reduced version of the N-TICK retains sensitivity to genetic-, environmental- and pharmacological-modulations of lifespan. The results show that N-rTICK is still sensitive to all the aforementioned differences, but shows larger and non-random errors. In particular, the pattern learned on the captive fish does not generalize well to the wild fish (dataset 5). In fact, the predicted age seems not to increase with time for the wild fish.

### 1.8 Heterogeneity of the GO term annotations of the N-TICK and N-rTICK

To understand better the composition of the 420 and 80 genes independently selected by N-TICK and N-rTICK, we first evaluated their intersection that comprises 29 genes and is statistically larger than what expected for random sets Fig. A13. This suggests that a small set of genes exisist that are good predictors of the aging process. The 15 human orthologs of these 29 genes are listed in Tab. 9. Its small size prevents the application of Gene Ontology Overrepresentation Analysis (ORA) in this set. Thus, We performed two ORAs on the 264 and 44 human orthologs of the genes selected by N-TICK and N-rTICK, respectively. In Tab.s 10 and 11, up to the first 10 most enriched annotations are listed for the genes selected by the N-TICK and N-rTICK, respectively. To visualize the relationships in complex lists of GO terms we used REVIGO [57]. The scatter plots of all the GO terms overrepresented in the two lists are reported in Fig.s 5 and 7.

**Fig. 5:**
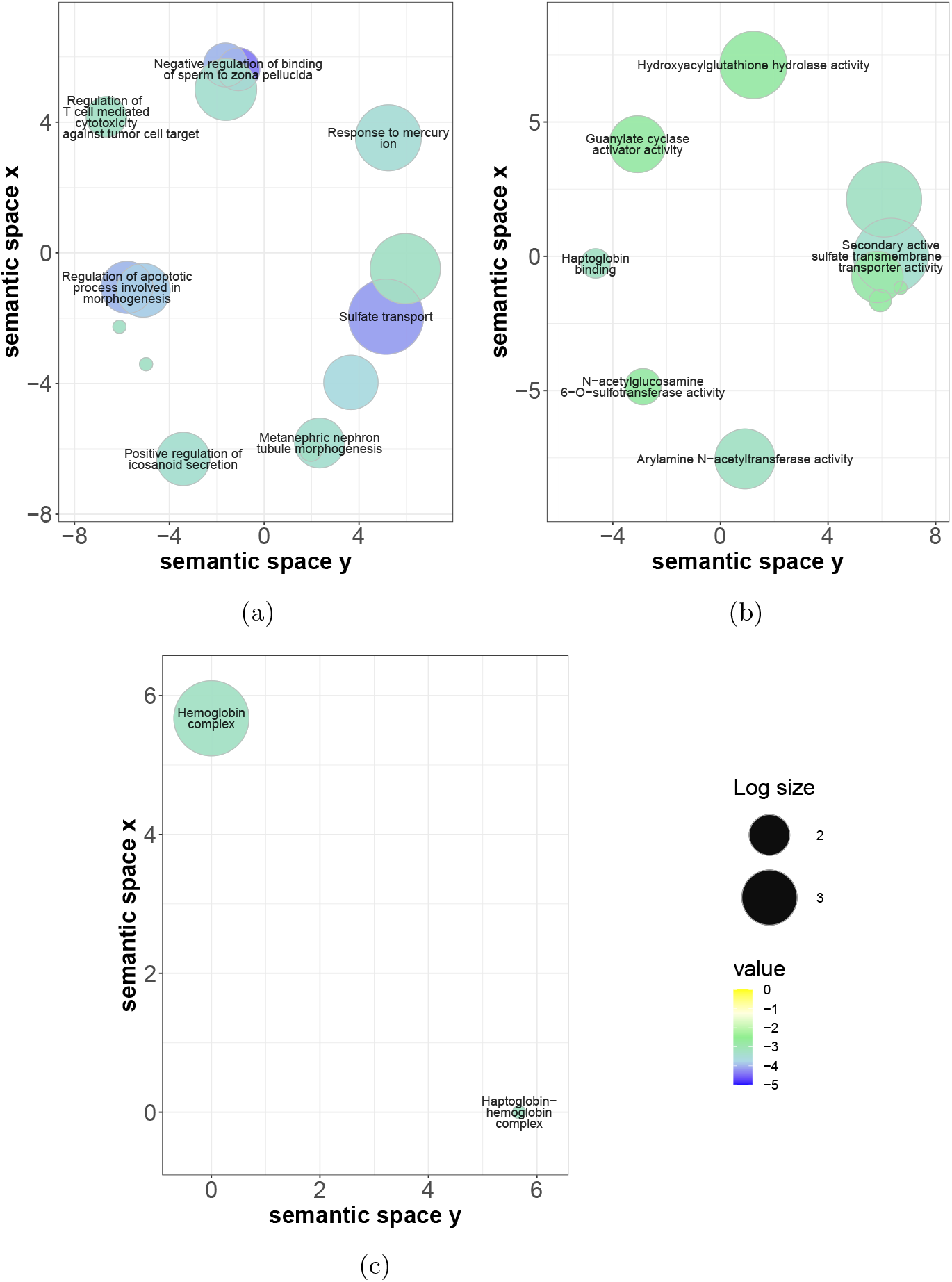
Revigo’s scatterplots of the gene set enrichment analysis of the 420 genes selected by our N-TICK. Fig. (a) visually summarizes the most relevant GO biological process, (b) the GO molecular functions, (c) the GO cellular components.

**Fig. 6:**
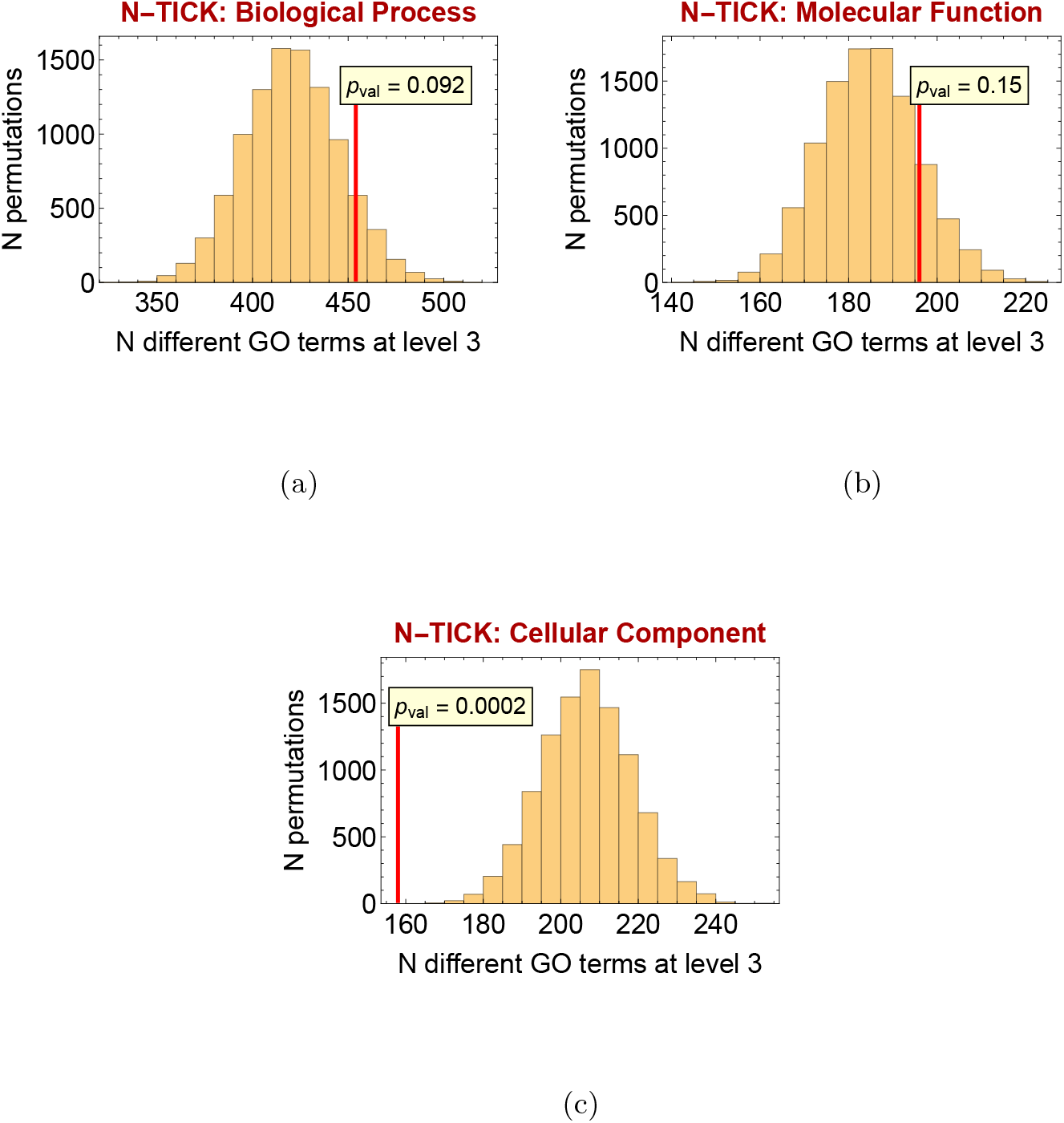
Permutation analysis to test the heterogeneity of the GO terms associated to the 420 genes selected by our N-TICK.

**Fig. 7:**
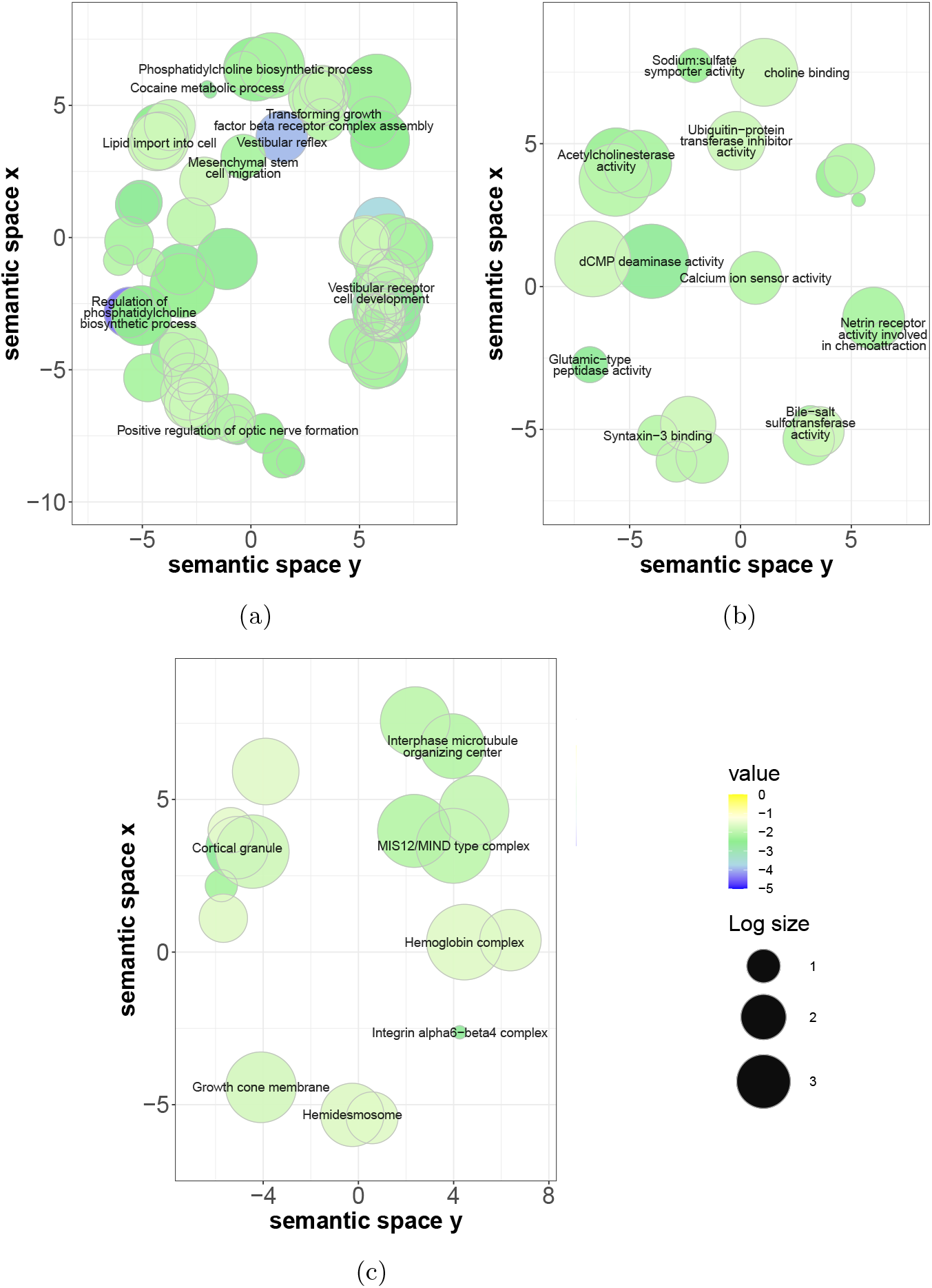
Revigo’s scatterplots of the gene set enrichment analysis of the 80 genes selected by our N-rTICK. Fig. (a) visually summarizes the most relevant GO biological process, (b) the GO molecular functions, (c) the GO cellular components.

**Table 9:** Known functions of the orthologs of the 15 genes in common between the ones selected by the N-TICK and N-rTICK. Source: [58].

| Gene name | Protein coding | Function |
| --- | --- | --- |
| ASTL | yes | It is involved in sperm and egg adhesion and fertilization. |
| DNAH3 | yes | It is involved in producing force for ciliary beating by using energy from ATP hydrolysis. It is also involved in sperm motility. |
| FMOD | yes | It may play a role in the assembly of extracellular matrix and in collagen fibrillogenesis. |
| HBG1 | yes | It encodes for one type of gamma chains that make up the fetal hemoglobin (HbF) which is normally replaced by adult hemoglobin (HbA) at birth. |
| HEPACAM2 | yes | It encodes a protein related to the immunoglobulin superfamily that plays a role in mitosis. |
| KRTCAP3 | yes | Supposedly, it encodes an integral component of membrane. |
| NTSR1 | yes | It mediates the multiple functions of neurotensin, such as hypotension, hyperglycemia, hypothermia, antinociception, and regulation of intestinal motility and secretion. |
| PAX2 | yes | It encodes for a transcription factor gene family that has a critical role in the development of the urogenital tract, the eyes, and the central nervous system. It may also have a role in kidney cell differentiation. |
| PPEF2 | yes | It encodes a member of the serine/threonine protein phosphatase with EF-hand motif family. Supposedly, it plays a role in the visual system in phototransduction. It may also function as a calcium sensing regulator of ionic currents, energy production or synaptic transmission. |
| SAG | yes | It belongs to the arrestin/beta-arrestin protein family and it is believed to participate in agonist-mediated desensitization of G-protein-coupled receptors and cause specific dampening of cellular responses to stimuli such as hormones, neurotransmitters, or sensory signals. It may play a role in preventing light-dependent degeneration of retinal photoreceptor cells. |
| SH2D7 | yes | An important paralog of this gene is SH2D2A, that encodes an adaptor protein thought to function in T-cell signal transduction. |
| SLC13A1 | yes | It encodes for an apical membrane Na(+)-sulfate cotransporter involved in sulfate homeostasis in the kidney. |
| SULT2A1 | yes | It encodes a member of the sulfotransferase family. Sulfotransferases aid in the metabolism of drugs and endogenous compounds by converting these substances into more hydrophilic water-soluble sulfate conjugates that can be easily excreted. SULT2A1 catalyzes the sulfation of steroids and bile acids in the liver and adrenal glands. |
| SYT1 | yes | It encodes for a member of the synaptotagmins which are integral membrane proteins of synaptic vesicles thought to serve as Ca(2+) sensors in the process of vesicular trafficking and exocytosis. It plays a role in dendrite formation by melanocytes. |
| ZBTB20 | yes | It encodes a transcription factor that plays a role in many processes including neurogenesis, glucose homeostasis, and postnatal growth. |

**Table 10:**
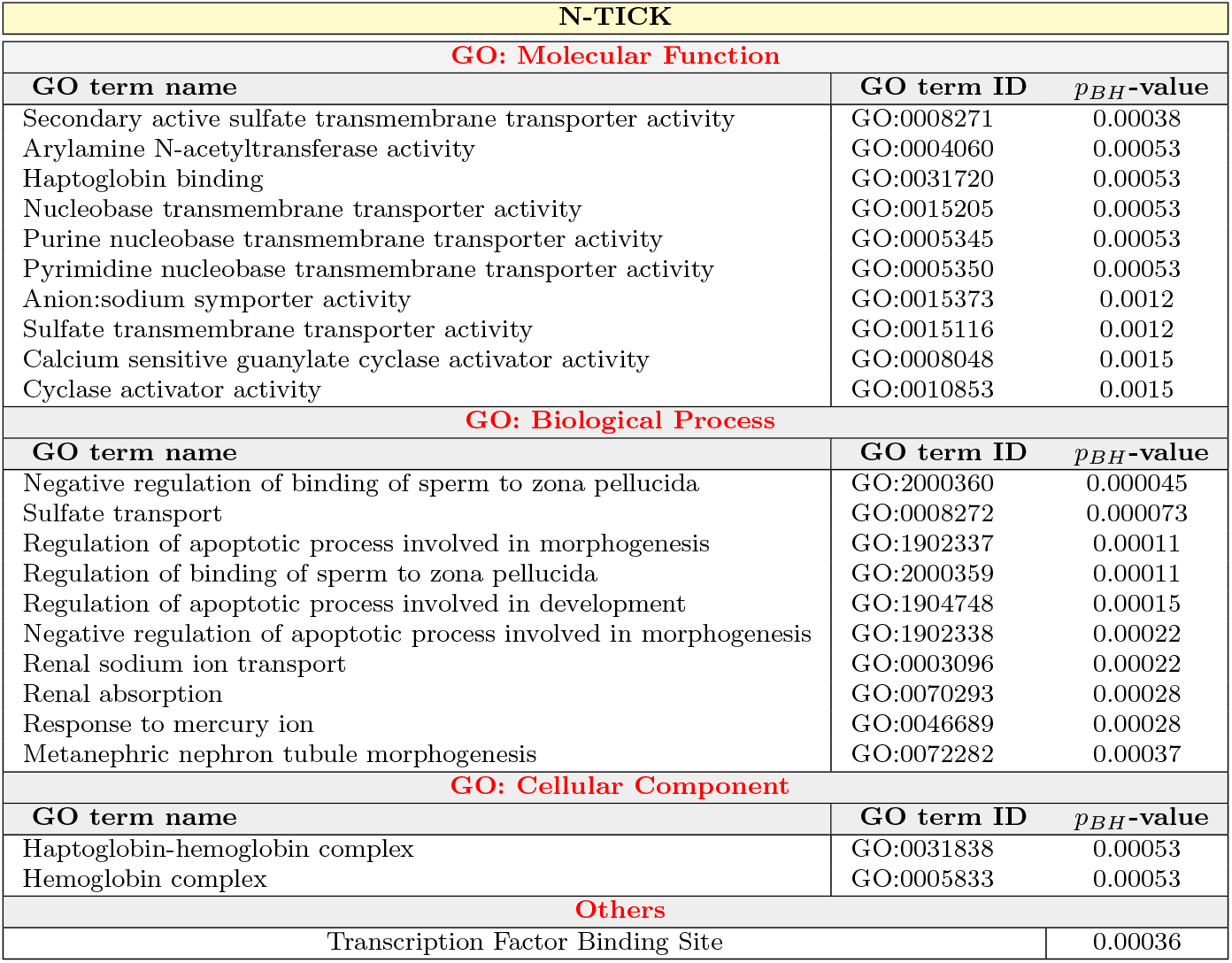
Summary of the gene set enrichment analysis for the 420 genes selected by our N-TICK. Up to the first 10 GO terms are listed in this table.

| N-TICK |  |  |
| --- | --- | --- |
| GO: Molecular Function |  |  |
| GO term name | GO term ID | $p_{BH}$-value |
| Secondary active sulfate transmembrane transporter activity | GO:0008271 | 0.00038 |
| Arylamine N-acetyltransferase activity | GO:0004060 | 0.00053 |
| Haptoglobin binding | GO:0031720 | 0.00053 |
| Nucleobase transmembrane transporter activity | GO:0015205 | 0.00053 |
| Purine nucleobase transmembrane transporter activity | GO:0005345 | 0.00053 |
| Pyrimidine nucleobase transmembrane transporter activity | GO:0005350 | 0.00053 |
| Anion:sodium symporter activity | GO:0015373 | 0.0012 |
| Sulfate transmembrane transporter activity | GO:0015116 | 0.0012 |
| Calcium sensitive guanylate cyclase activator activity | GO:0008048 | 0.0015 |
| Cyclase activator activity | GO:0010853 | 0.0015 |
| GO: Biological Process |  |  |
| GO term name | GO term ID | $p_{BH}$-value |
| Negative regulation of binding of sperm to zona pellucida | GO:2000360 | 0.000045 |
| Sulfate transport | GO:0008272 | 0.000073 |
| Regulation of apoptotic process involved in morphogenesis | GO:1902337 | 0.00011 |
| Regulation of binding of sperm to zona pellucida | GO:2000359 | 0.00011 |
| Regulation of apoptotic process involved in development | GO:1904748 | 0.00015 |
| Negative regulation of apoptotic process involved in morphogenesis | GO:1902338 | 0.00022 |
| Renal sodium ion transport | GO:0003096 | 0.00022 |
| Renal absorption | GO:0070293 | 0.00028 |
| Response to mercury ion | GO:0046689 | 0.00028 |
| Metanephric nephron tubule morphogenesis | GO:0072282 | 0.00037 |
| GO: Cellular Component |  |  |
| GO term name | GO term ID | $p_{BH}$-value |
| Haptoglobin-hemoglobin complex | GO:0031838 | 0.00053 |
| Hemoglobin complex | GO:0005833 | 0.00053 |
| Others |  |  |
| Transcription Factor Binding Site |  | 0.00036 |

**Table 11:**
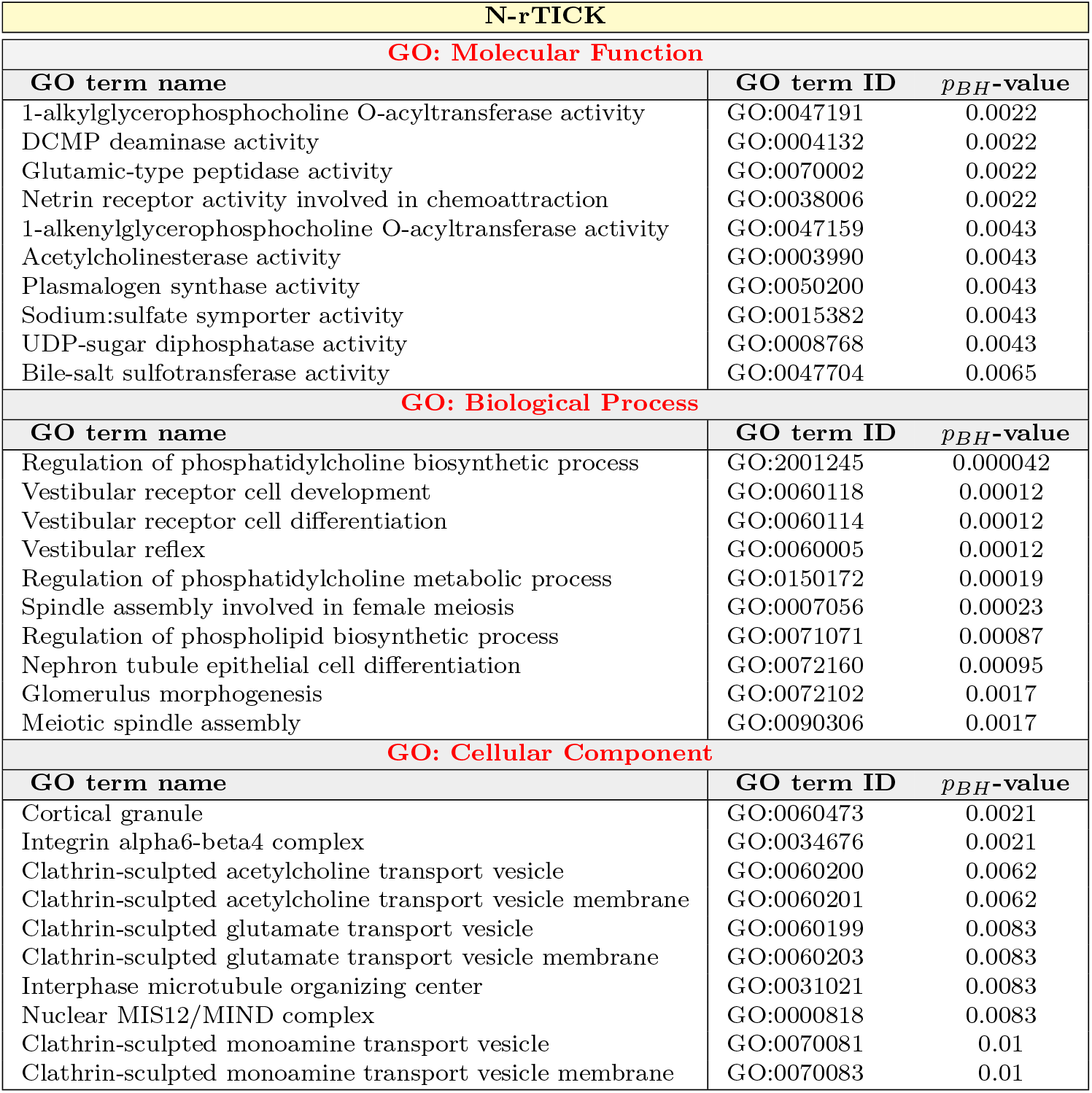
Summary of the gene set enrichment analysis for the 80 genes selected by our N-rTICK. Up to the first 10 GO terms are listed in this table.

The most striking aspect of this analysis is the large number and wide heterogeneity of significantly overrepresented GO terms in the two lists, which is particularly noticeable for N-rTICK. Thus, we performed a permutation test to quantify the diversity of GO terms and assess if the number of GO terms associated to the genes selected by the two clocks is higher than what is detected by sampling an equal number of randomly selected genes. Given that the leaves of the GO tree contain hyperspecific descriptors, that can be traced back to a common pathway, we decided to cut the GO tree at level 3, meaning that any GO term found at level 4 or deeper was converted to its ancestor at level 3. The results of the permutation tests obtained by counting the GO terms in this way are illustrated in Fig. 6 and 8 for our N-TICK and N-rTICK, respectively. As it can be noted, the number of the GO terms associated to the genes selected by the two clocks is higher than the average of the permutation distributions in the ”Biological Process” and ”Molecular Function” domains and this difference is significant for N-rTICK. The number of GO terms in the ”Cellular Component” domain, instead, is significantly lower for the N-TICK and perfectly in line with a random selection for the N-rTICK. In all the three cases, the N-rTICK genes are more heterogeneus than those selected by the N-TICK. This result indicates that an accurate prediction of age from a expression of a limited number of genes requires sampling diverse biological processes.

**Fig. 8:**
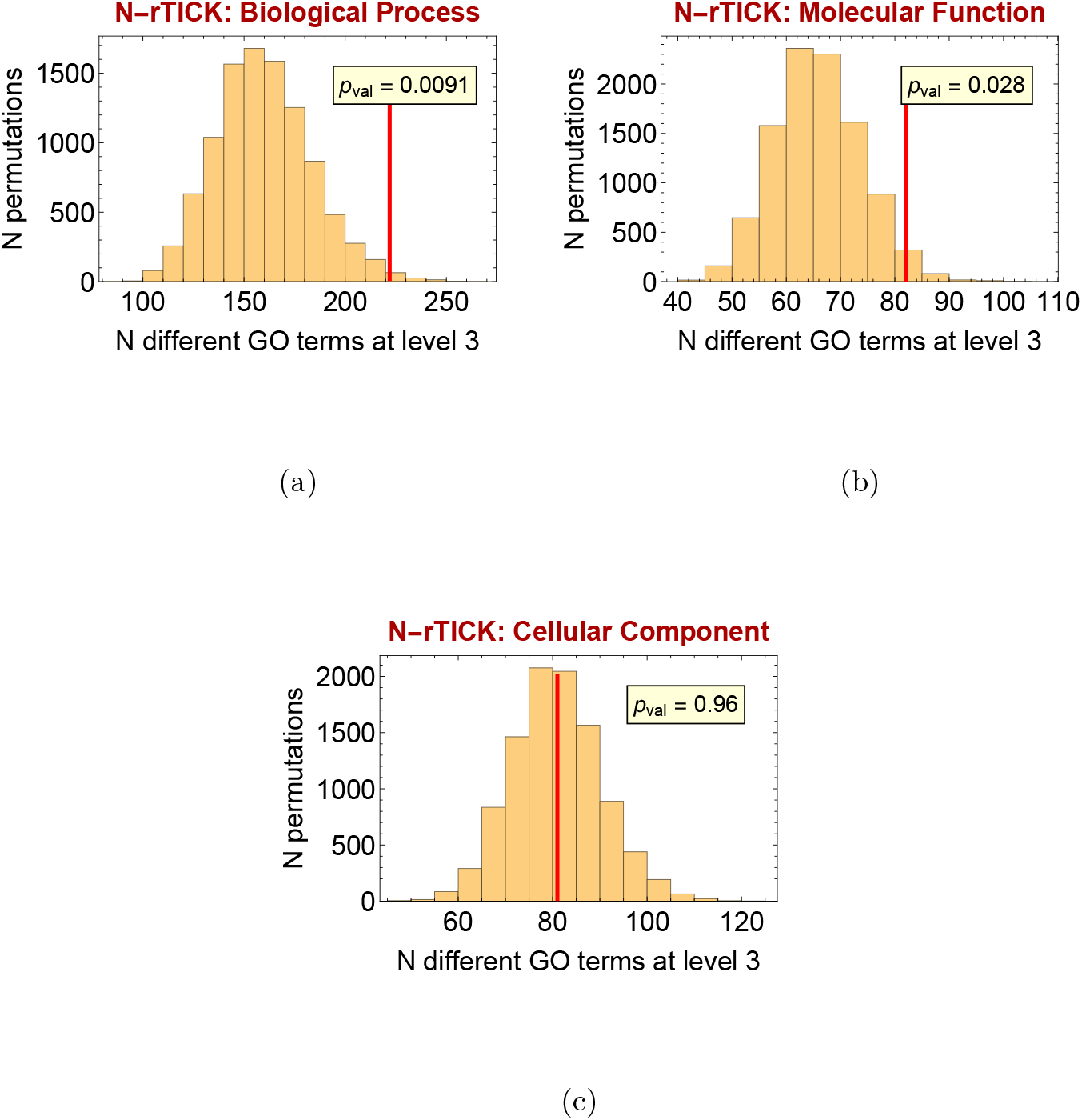
Permutation analysis to test the heterogeneity of the GO terms associated to the 80 genes selected by our N-rTICK.

### 1.9 The genes selected by N-TICK and N-rTICK provide a multi-tissue age predictor in humans

To assess the translability of our *N. furzeri* -based clocks to humans, we tested if the human orthologs of the genes selected by the latters can be used also to predict human age. Thus, we modified the SA-DNN to receive as input the expression levels of the 293 human orthologs for the genes selected by the N-TICK and N-rTICK (see Fig. A13b) and we deactivated its selector layer. The dataset used for this application is the public GTEx dataset (dataset 6), in which age is discretized into classes spanning one decade. Thus, the architecture described so far was trained to develop two different types of predictors: a classifier that outputs an age class and a regressor trained to return the middle age of the decade. We refer to these different models as H-cTICK and H-TICK, respectively. In both of them, the adversarial learning framework was used to minimize the influence on prediction of the following confounders: tissue, sex, type of death and subject identifier.

Fig. 9a and 10a show the performance of the H-cTICK and H-TICK, respectively, computed on the training, validation and test sets and on an hold-out set composed solely by liver samples: a tissue that is not included in any of the other sets. As it can be seen, in both the cases the performance is very high and stable across the different sets. Note that in the regressor application, the age label has an inherent discretization error that, supposing uniform the age distribution, would generate an RMSE of 2.87 and a MAE of 2.5. This means that our predictions are almost as accurate as possible.

**Fig. 9:**
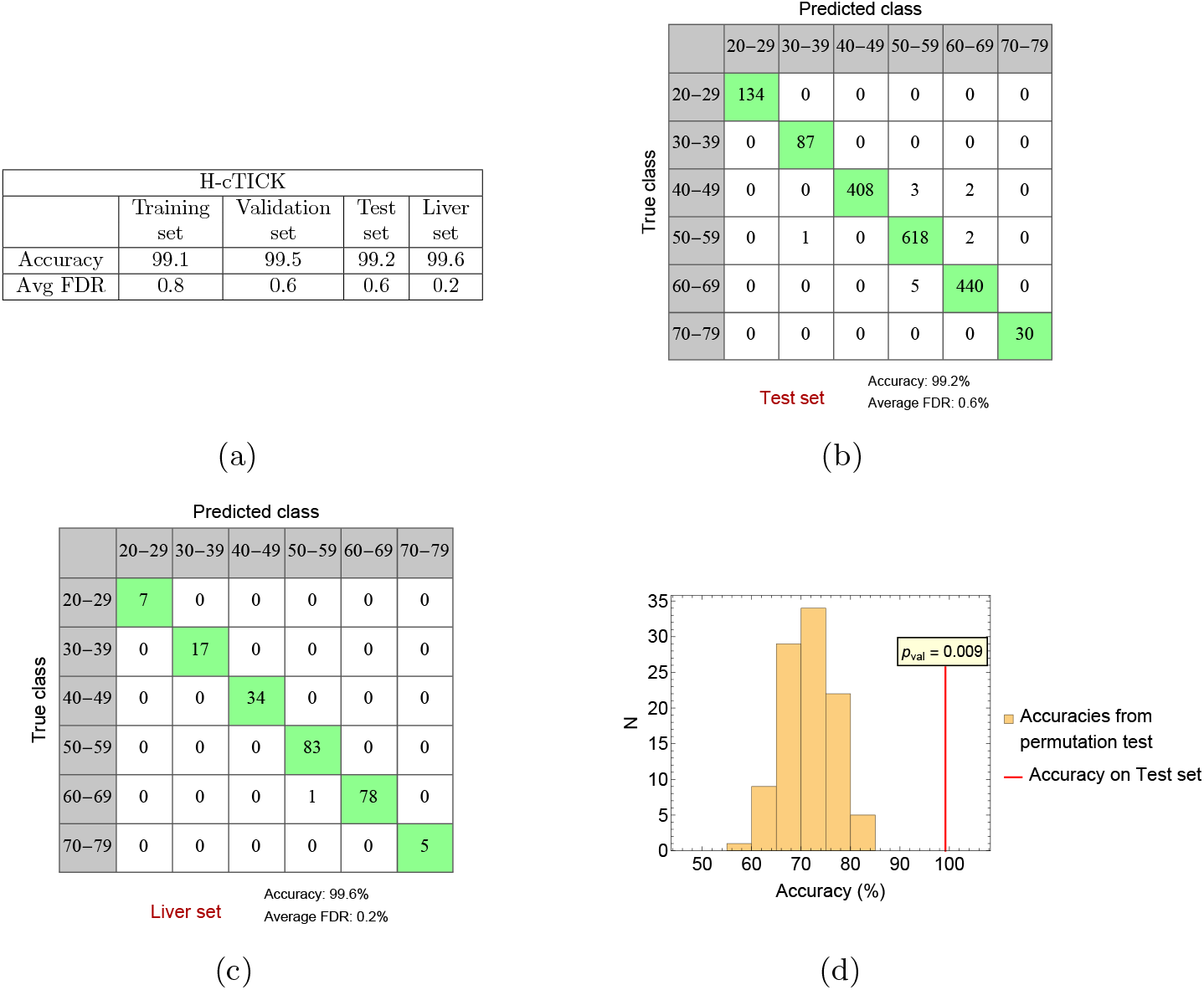
Performance of the H-cTICK clock. Fig. (a) shows the accuracy and the FDR computed on the training, validation, test sets and an hold-out set composed solely by a tissue not represented in the previous sets: the liver. Fig.s (b) and (c) are the confusion matrices computed on the test and the liver hold-out sets, respectively. Fig. (d) illustrates the permutation test performed to verify if the genes selected by the N-TICK and N-rTICK allow to reach an higher accuracy rather than using random lists of genes of equal length.

**Fig. 10:**
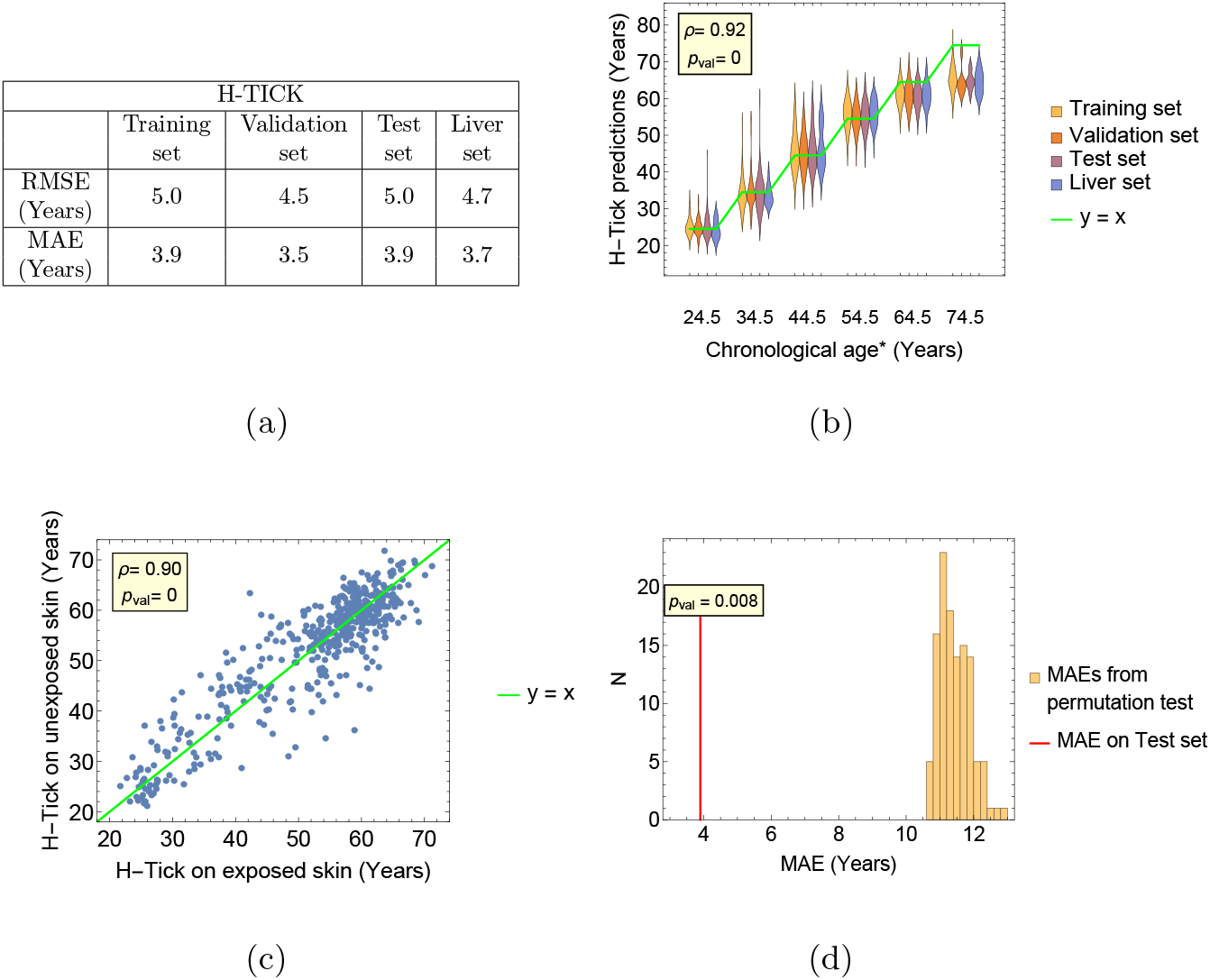
Performance of the H-TICK clock. Fig. (a) reports the RMSE and the MAE computed on the training, validation, test sets and an hold-out set composed solely by a tissue not represented in the previous sets: the liver. Fig. (b) is a violin plot showing the predicted vs the true age of all the sets. Fig. (c) is a scatter plot of age prediction on exposed vs unexposed skin samples. Fig. (d) illustrates the permutation test performed to verify if the genes selected by the N-TICK and N-rTICK allow to reach a lower MAE rather than using random lists of genes of equal length.

**Fig. 11:**
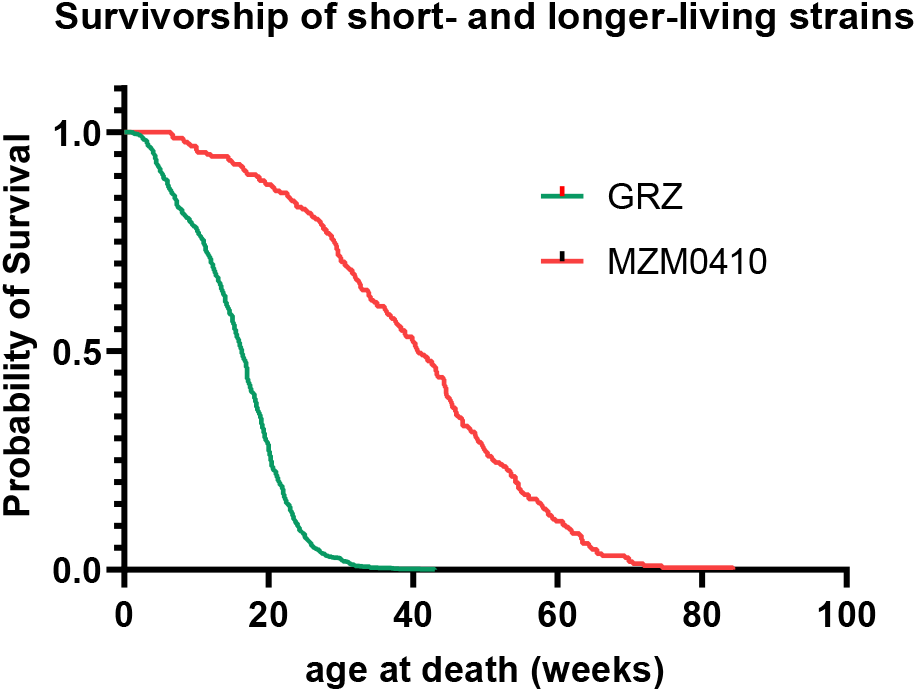
Survival curves of GRZ and MZM0410 *N. Furzeri* ’s strains in the facility that collected the data of this study.

In Fig. 10b, it can be noted that the age of the 7th decade seems underestimated. However, according to the general description reported on the GTEx website, the subjects included in the study are in the age range of 20-71 years. If this information is correct and updated to the latest release, the underestimation observed in the 7th decade is not only correct but even remarkable. In fact, despite the algorithm has been trained to predict the mean value of each decade, it is able to detect the true age. Notably, the H-cTICK perfectly predicts the 7th decade even considering that this set of data spans only 2 years of age (Fig. 9b-c).

An analysis reported in Sec. A.3 shows that the performance of the H-cTICK and the H-TICK are almost invariant across different subcategories of the dataset, defined by both the variables for which the clocks were made resilient through adversarial learning and other ones. The high generalizability achievable using our SA-DNN is also confirmed by the H-TICK insensitivity to photoaging, despite the fact that exposed and unexposed skin samples were not treated as different instances of the tissue confounder. In fact, Fig. 10c shows that the H-TICK predicted age of the exposed and unexposed skin samples belonging to the same patients are highly correlated with a symmetrical distribution with respect to the *y* = *x* line.

Finally, we performed a permutation test to verify if the genes selected with our N-TICK and N-rTICK are significantly more informative on the aging process of humans than other sets of an equal number of randomly selected genes. As it can be seen form Fig. 9d and 10d, the genes selected on the *N. Furzeri* allows the H-cTICK to reach a significantly higher accuracy and the H-TICK a significantly lower MAE, with respect to any of the random set of genes tested.

## 2 Discussion

In this study, we have developed a multi-tissue transcriptomic clock for *N. furzeri*, that we named N-TICK, whose performance is close to the theoretical limit. It is interesting to compare the precision of N-TICK with that of other molecular aging clocks, even with the limitations that: errors are measured using different metrics, survivorship curves and phenotypic variability can significantly differ across different species. The *N.furzeri* strains of datasets 1 and 2 have a median lifespan of ∼ 8 months and humans have a median lifespan of ∼ 80 years. If we scale the errors linearly with the lifespan, N-TICK prediction errors would be comparable to an error of 2-4 months in the age prediction of humans, almost one order of magnitude smaller than the minimum error computed on humans, that is of 3 years [44] (see Table 8). The last two columns of Table 8 report the measured error of each clock and the age span of the samples used in the study. Although a fair comparison is not possible for the above reasons, if we use error to age span ratio as a metric, N-TICK reaches the lowest error of 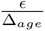 = 0.0038, outperforming all the other clocks. This result, as well as those of others [17, 59], demonstrates that gene expression data contain sufficient information for accurate age prediction rivalling DNA-methylation data and that the noise originated by the dynamic nature of this type of data can be overcome by using our SA-DNN method.

Notably, the errors of N-TICK are randomly distributed with respect to age. Instead, many clocks based on elastic net regression display a clear age over-estimation/underestimation of youngest/oldest individuals [60, 61]. In our opinion, this phenomenon of ”non-linear rate of clock ticking” [22], which is considered a property of aging clocks deriving from the basic biology of aging, should be instead interpreted as a limitation of linear models in modelling non-linear aging trajectories. The use of a DNN, instead, allows to incorporate the non-linearity of aging into the model, making the error of the predictions independent from the individual’s age. Indeed, the reduced version of N-TICK, N-rTICK, that uses only 80 genes as input, provides predictions with an error to age span ratio of 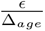 = 0.038, showing that reducing the number of input increases the errors, and these errors are non-randomly distributed. Therefore, reduced prediction accuracy at the extremes of the tested age range is not a general property of clocks, but an indicator that the predictor has not reached the theoretical limit. In this context, it is worth stressing that in the application of the N-TICK to the wild brain samples, we obtained overestimation of the predicted age but also a slope that, with respect to the true one, is inferior to 1. It should be noted that wild fish reach asymptotic size much earlier than captive ones [56] and in the age range explored, the former ones do not show growth [55]. Although three points in lifespan are not sufficient to reliably interpret this phenomenon, we can speculate that the N-TICK predictions correctly capture the heterochronic and non-linear nature of growth and aging in the wild vs. captivity.

The broad interest in the development of aging clocks derives from their possible use as biomarkers in intervention studies [23] and the fact that they capture the effects of life-span modulating interventions in experimental organisms [15–17] and to some extent in humans [11, 12]. N-TICK is sensitive to genetic-, environmental- and pharmacological-modulations of lifespan. In particular, the conditions under which we observed an overestimation/underestimation of the predicted age are those in which life extension/reduction were observed experimentally. *N. furzeri* is a convenient model organism to investigate the effects of experimental intervention on aging and lifespan [18]. N-TICK can be used as a biomarker in pilot experiments using a set of compounds to select those that may have life-extending effect and deserve further investigation. Moreover, the fact that the same gene set that constitutes the input of N-TICK is also able to provide an accurate age estimation in humans is of particularly high relevance in a context of intervention testing as it ensures a translability of the results.

The analyses on the N-rTICK show that an accurate age prediction with an error to age span ratio of 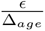 = 0.038 can be obtained analyzing the expression levels of only 80 genes. However, reducing the number of input data not only increases the errors, it also makes them non-randomly distributed and it makes the predictor less generalizable to other data, as explained in Sec. 1.7.

The assumption that the clock’s prediction error are not solely due to random errors, but contain information on the ”bioloigcal age” of the individual is key in the application of aging clocks. If the transcriptome contains sufficient information to predict anagraphic age with an almost perfect accuracy, and the errors occur because of the tug-of-war between accuracy and the number of input selected, how to choose the right threshold leading to a correct estimation of the biological age? The patterns in the observed N-rTICK’s errors, i.e. their non-random distribution and their higher values for data collected under different conditions, suggest that the errors do not solely depend on the different aging speed of an individual, but also on other factors such as the age itself and the similarity between the data used to train the algorithm and those to which the algorithm is applied. Indeed, there is still an open debate as to whether deviation from the anagraphic age detected by epigenetic clocks contain information on individual risk. For instance, a recent analysis of five epigenetic clocks on over 1000 participants did not detect association of higher epigenetic age with functional decline [62]. Thus, in our opinion, the use of aging clocks should be limited to two applications:

1. to anticipate results of intereventions, as detailed above, if they are trained to minimize the prediction error and they prove to be resilient to over-fitting and reliable on a wide heterogeneous cohort of individuals. In this case the gene filtering should be exclusively used to avoid overfitting and not to minimize the data used by the clock.
2. to understand the composition of the genes that better recapitulate the aging process. In this case, heavily filtering the input data is worth pursuing. In fact, by increasing the penalty in the filtering layer, the model will select only one gene that best approximates the contribution to the age estimation of a set of highly correlated genes that carry similar information.

As a consequence of the second point above, an enrichment of specific pathways should not be prominent, because the gene selection process will aim towards sampling as many pathways as possible, with as few genes as possible. As a confirmation, our permutation analysis has shown that the biological GO terms associated to the genes selected by our N-rTICK, where a heavy filtering is imposed, are more heterogeneous than those associated to a random gene set of equal size. This also may indicate that the information necessary to infer age accurately is distributed on genes involved in diverse pathways and is not concentrated in a small set of ”master genes” belonging to a few biological processes. This observation seems to support the deleteriome hypothesis, claiming that the aging process arises from a pletora of different detrimental alterations to cellular and organismal function accumulating over time [63].

We took advantage of the orthology of the genes constituting the *N. furzeri* clocks to develop a reliable human clock based on the same genes that could provide a nearly-perfect classification of human samples into decade bins. Furthermore, when we performed a regression analysis, this set of human genes provided an age prediction whose error that is on the average 2.5 times smaller than a randomly selected gene set of equal size. This and previous results [64, 65] indicate that the transcriptomic correlates of the aging process are well conserved across these two vertebrate species and can be described by the expression level of a limited number of genes, further confirming that *N. furzeri* is a valid animal organism to model human aging. On the other hand, the selected genes are not necessarily the best possible set to accurately predict age of human samples. In our opinion, the N-TICK and N-rTICK selected a list of genes that effectively summarizes the pathways involved in the aging process in *N. furzeri* but it is entirely possible that a different set may provide higher accuracy. The permutation test shows that it is very unlikely to find such an alternative and heterogeneous set by random sampling. Interstingly, however, even a random gene set can be used to provide a prediction with a MAE of 12.5 year. This results indicate how effectively the SA-DNN can identify age-related patterns of gene expression and also suggest that the human transcriptome contains diffuse information on age-dependent processes [13].

The predictions of the H-TICK and H-cTICK models are almost as accurate as possible, considering that the age labels used to train the algorithm are discretized into decades. As explained in Sec. 1.9, the apparent age underestimation of our H-TICK on the 7th decade is a positive point because this decade is populated by subjects of 70-71 years. Furthermore, the fact that the H-cTICK perfectly predicts the 7th decade indicates instead that these clocks may have a higher precision and accuracy with respect to what we can measure with the age labels available.

Finally, the H-TICK age predictions do not show any systematic overestimation of the exposed skin samples with respect to the unexposed ones, further confirming the high generalizability of the learning model adopted. We interpret this result as an indirect effect of the adversarial learning framework, that has forced the model to learn a pattern that predicts age and is invariant in the different tissues. Therefore, a pattern that differentiates sun-exposed and non-exposed skin was penalized.

## 3 Conclusion

We conclude that our SA-DNN architecture has proved to be resilient to both overfitting and confounding effects across all the analyses presented in this paper. Furthermore, its predictions correctly detect experimental conditions that modulate lifespan and the genes selected in *N. furzeri* provide a clock applicable on human data. Finally, thanks to its tunable selector layer, SA-DNN can vary the trade-off between accuracy and number of input variables, an issue particularly relevant for the design of scalable assays. Our SA-DNN represents the prototype of a new class of aging clocks that provide biomarkers applicable to intervention studies in model organisms and humans and we recommend the use of SA-DNN or similar architectures for further studies on biological clocks.

## 4 Methods

### 4.1 *N. furzeri* ’s samples preparation

Fish were bred at the Leibniz Institute on Aging, Fritz Lipmann Institute, Jena. The survival curves of the GRZ and MZM0410 strains measured in our facility without any intervention are reported in Fig.11. Procedures for fish breeding, husbandry and euthanasia were performed in accordance with the rules of the German Animal Welfare Law and approved by the Landesamt für Verbraucherschutz Thüringen, Germany. Details on husbandry procedures are detailed elsewhere [37]. At the desired age, fish were sacrificed via anesthetic overdose (Tricaine, MS-222), in accordance with the prescription of the European (Directive 2010/63/UE) and Italian law (DL 26/04-03-2014), and the tissue was immediately frozen. RNA extraction followed described procedures [37].

### 4.2 RNA sequencing of Nothobranchius’s samples

In general, sequencing of RNA samples was performed using Illumina’s next-generation sequencing methodology [66]. In detail, total RNA was quantified and quality checked using Agilent 2100 Bioanalyzer Instrument (RNA 6000 Nano assay). Libraries were prepared from 500 ng of total RNA using TruSeq RNA v2 library preparation kit (Illumina) according the manufacturer’s instructions and subsequently quantified and quality checked using Agilent 2100 Bioanalyzer Instrument (DNA 7500 assay). Libraries were sequenced using a HiSeq 2500 System running in 51 cycle/single-end/high output mode. Sequence information was converted to FASTQ format using bcl2fastq v1.8.4. The RNA sequencing reads were aligned to the *N. furzeri* genome assembly (assembly version Nfu 20150522, gene annotation version 150922, both downloaded from the Nothobranchius furzeri Information Network Genome Browser) [67] using STAR 2.5.1b (parameters: –alignIntronMax 100000, – outSJfilterReads Unique) [68]. For each gene, all reads that map uniquely to one genomic position were counted with FeatureCounts 1.5.0 under default settings [69]. Sequencing, mapping and counting qualities were assessed with MultiQC 0.9 [70].

### 4.3 Preprocessing and normalization of the *N. furzeri* ’s datasets and the GTEx dataset

For each of the five *N. furzeri* ’s datasets and for the GTEx dataset the following operations have been apllied to the data:

- the genes with a length inferior to 500 bp have been discarded, since count estimation for short genes is less reliable.
- The genes with less than 10 counts in at least 20% of the samples belonging to the same tissue have been filtered out.
- For multi-tissue datasets, the intersection of the gene lists selected for each tissue, according to the previous operations, has been selected.
- The raw-counts filtered, according to the previous points, were normalized with the ”gene length corrected trimmed mean of M-values” (GeTMM) algorithm [71], that corrects for sequencing depth, gene length and total sample RNA output.
- Finally, the data were Log2 transformed.

All datasets were preprocessed separately following the steps described above. Additionally, when a dataset is split into multiple subsets (like training, validation, test and hold-out sets) each of them is normalized with the GeTMM individually, to avoid information leakage from one set to the other.

### 4.4 Our Selective Adversarial Deep Neural Network approach: SA-DNN

Our SA-DNN algorithm was designed to specifically address three problems that often affect biomedical applications based on deep learning: (i) the overfitting phenomenon [72], (ii) the confounding effects [35] and (iii) the difficulty to access the pattern learned by the algorithm [73].

Our SA-DNN addresses the overfitting problem using dropout layers [74] and implementing a new filtering regularized layer, that also increases the intrepretability of the model, facing also the third issue. The detrimental effects of confounders on the learning process is solved by our SA-DNN by the adoption of an adversarial learning framework [75].

In the next sections, we are going to describe the two distinctive features of our algorithm: the filtering layer and the adversarial learning framework and how they are combined in the SA-DNN architecture used to develop our clocks.

#### 4.4.1 Filtering layer

In addition to the adoption of dropout layers, to reduce overfitting we propose the use of a new filtering layer as the first layer of our deep architecture. We call it the *selector* layer. This is a simple binary layer that multiplies the input features by either 0 or 1, where the multiplication by 0 eliminates the corresponding feature from consideration by subsequent layers. To force the filtering layer to multiply as many features as possible by 0, we use a L1 regularization on the sum of its weights. In this way, the network is not only trained to minimize the prediction error, but it is also penalized whenever it assigns a non-zero weight to a feature. This means that the network is encouraged to always assign 0, unless that feature is so important that not using it introduces large errors in the prediction.

Note that binary weights are not differentiable and thus cannot be trained with standard backpropagation algorithms based on gradient descent, because the gradient cannot be calculated for discrete weights. To solve this issue, we decided to use continuous weights constrained between 0 and 1 during the gradient calculation, and then binarize them by rounding them to the nearest integer at prediction time. With this strategy however, we empirically noticed that during long training sessions, most weights were pushed below 0.5 and would not be able to go back up, as the gradient calculation is not aware of the rounding. To mitigate this problem we modified the L1 regularization introducing a threshold *t*, a minimum absolute sum of weights below which the regularization would be inactivated, as shown in the following formula:

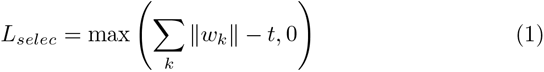

As a consequence the choice of the threshold *t* can influence the number of feature with a non-zero weight. As a rule of thumb, small/large values of *t* selects a small/large number of inputs to use for the prediction.

#### 4.4.2 Adversarial learning

Our approach to solve the problem of confounders involves the use of adversarial learning (AL). AL is a technique that consists in a two-players game between two modules that perform two different tasks, in which each module is iteratively trained to maximize its performance possibly at the expenses of the other module. In our application, the two competitive tasks are the age prediction and the prediction of a list of confounders that we want our clock to be resilient from. The implementation of our adversarial learning framework is inspired by [75], that uses a tripartite structure composed by a module that extract a condensed representation of the input data, and two modules that perform the two competitive tasks starting from the condensed representation.

#### 4.4.3 The architecture of our SA-DNN used to develop our clocks: N-TICK, N-rTICK, H-TICK and H-cTICK

Formally, the development of our clocks can be described as a regression problem for which each of the *N* instances *i* is a triplet, composed by a vector of input features, the expected output and a vector of confounders (potentially containing both categorical and continuous variables): 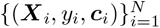. We aim to predict *y_i_* from a set of features ***F****_i_* derived from a subset of the input ***X****_i_* ensuring that ***F****_i_* does not contain information about the confounders ***c****_i_*. We achieve this objective using the DNN illustrated in Fig. 12. The net is composed basically by 3 parts:

- The Feature extractor FE The FE processes the ***X****_i_* to extract ***F****_i_*. The FE is composed by an initial *selector* layer (described in the previous section), followed by fully connected layers alternated to dropout ones. In Fig. 12 the symbol *θ*_FE_ indicates all the parameters of the FE.
- The Predictor ℙ P takes as input the ***F****_i_*, generated by the FE, and tries to predict the target output *y_i_*, producing an estimated *ŷ_i_*. The structure of ℙ is composed by fully connected layers and its parameters are indicated with *θ*_ℙ_.
- The confounders predictor ℂ Similarly to ℙ, ℂ is composed by fully connected layers and tries to predict the counfounders ***c****i* from ***F****_i_*. Its parameters are indicated with *θ*_ℂ_ During the training process, the parameters of the DNN *θ*_FE_, *θ*_ℙ_ and *θ*_ℂ_ are adjusted to minimize the value of a cost function computed on a subset *B* of the available data, called batch. The parameters update happens in three rounds (see Fig. 12) and involves three different loss functions and the regularization *L_selec_* described before. Here we list the losses:
- A loss that minimizes the sum of the squared errors of the predictions made on the batch:

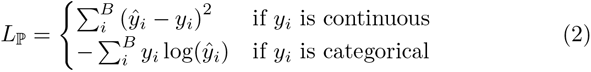

This loss is used to select the parameters that allow to correctly predict the task of interest. In our N-TICK, N-rTICK and H-TICK clocks *y_i_* is the age of the individual represented as a continuos variable, while for our H-cTICK *y_i_* is a categorical variable indicating the age class (spanning 10 years) of the individual.
- A loss that minimizes the sum of the errors made in predicting the value of the confounders *c_ij_*, where *i* and *j* are the indexes of the specific instance and confounder, respectively. The errors are computed differently for continuous confounder (using the root summed squared error) and categorical ones (using the cross-entropy)^1^:

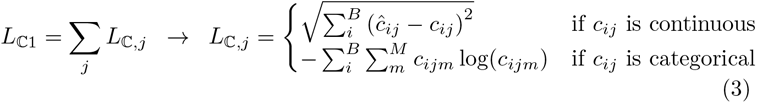

This loss is used for training the confounder predictor ℂ. The list of confounders corrected by each of our clocks are described in Sec. 1.2.
- A loss that minimizes the sum across all the *j* counfounders of the correlation between the vectors of the true and the predicted *j*-th confounder across the batch *B*.

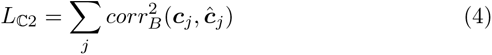

This loss is used to remove the influence of the confounder variables from the vector ***F****_i_*.

**Fig. 12:**
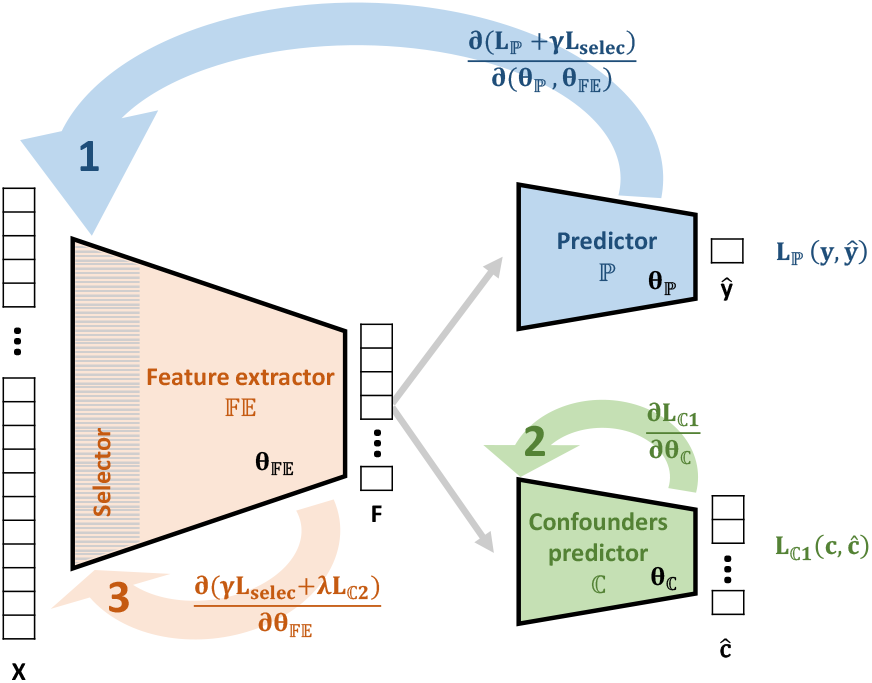
Graphical representation of the SA-DNN architecture developed and used in this work.

The adversarial approach is clearly visible in the last two losses, that perform conflicting tasks. In fact, while *L*_ℂ1_ tries to correctly predict confounders, *L*_ℂ2_ tries to modify ***F_i_*** so that it becomes impossible to do so. These last two losses also use different metrics. *L*_ℂ1_ uses the standard losses for continuous and categorical variables, to ensure that the output of ℂ is as close as possible to the confounder values. This in turn means that when using *L*_ℂ2_ to remove the dependency of the confounders from ***F****_i_* using a correlation metric, both linear and non-linear dependencies between ***F****_i_* and **c***_i_* are destroyed.

The training process iterates through the losses and starts with the of *θ*_FE_ and *θ*_ℙ_ according to *L*_ℙ_+ *γL_selec_*, where *γ* is a parameter that sets the strength of the filtering regularization. Then, the parameters *θ*_ℂ_ are updated according to *L*_ℂ1_, while keeping *θ*_FE_ (and thus ***F****_i_*) stationary. Finally, *θ*_FE_ is updated again to remove the effects of confounders, using the loss *γL_selec_*+ *λL*_ℂ2_, where *λ* is an hyperparameter that sets the strength of the adversarial learning. These three steps, repeated for every batch, are equivalent to optimizing *θ*_FE_ with a saddle-point loss of the form:

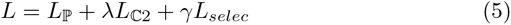

Note that *λ* is just mathematical representation of the combined effect of the different learning rates applied to the three steps.

### 4.5 Clocks training and validation scheme

#### 4.5.1 N-Tick and N-rTick

Both the N-TICK and N-rTICK were trained, validated and tested on the same sets of data. The parameter optimization was done training the algorithms on 180 samples taken from dataset 1, using a grid search exploring the parameter space outlined in Table 12. The hyperparameter selection was based on the comparison of the performance computed on the validation set, composed by 65 samples taken from dataset 1, with no intersection with the training set. The remaining 63 samples of dataset 1 were used as a test set as well as datasets 2-5. Note that the training, validation and test sets were fishwise split.

**Table 12:** Parameter space explored for the clocks developed on the *N. furzeri* data.

| Parameter | Range | N-TICK | N-rTICK |
| --- | --- | --- | --- |
| Neurons first layer ($\mathbb{F}\mathbb{E}$) | 30 – 100 | 64 | 64 |
| Total number of parameters | 10,000 – 500,000 | 60,000 | 15,000 |
| Hidden layers ($\mathbb{F}\mathbb{E}$) | 2 – 6 | 4 | 4 |
| Hiddern layers ($\mathbb{C}$) | 2 – 6 | 3 | 3 |
| Hidden layers ($\mathbb{P}$) | 2 – 6 | 3 | 3 |
| Dropout rate | 0.01 – 0.30 | 0.05 | 0.05 |
| Number of learned features ($\mathbb{F}\mathbb{E}$) | 10 – 100 | 30 | 30 |
| Learning rate ($\mathbb{F}\mathbb{E}$) | $10^{-9}$ – $10^{-4}$ | $2.5 \cdot 10^{-6}$ | $1 \cdot 10^{-7}$ |
| Learning rate ($\mathbb{C}$) | $10^{-9}$ – $10^{-4}$ | $1 \cdot 10^{-7}$ | $1 \cdot 10^{-7}$ |
| Learning rate ($\mathbb{F}\mathbb{E}+\mathbb{P}$) | $10^{-9}$ – $10^{-4}$ | $5 \cdot 10^{-6}$ | $5 \cdot 10^{-6}$ |
| $\gamma$ | $10^{-9}$ – $10^{-4}$ | $1 \cdot 10^{-6}$ | $5 \cdot 10^{-5}$ |
| $t$ | 100 – 10000 | 6000 | 500 |
| Batch size | 10 – 100 | 50 | 50 |

The threshold *t* and the learning rates of FE, FE + ℙ and ℂ are the only hyperparameters that were not chosen exclusively to maximize the performance on the validation set. In fact, *t* was chosen trying to find the best compromise between the performance and the amount of genes that we wanted our clock to use. The learning rates were chosen trying to remove all the information related to the confounders without sacrificing too much the age prediction. The final parameters of the clocks are reported in the last two columns of Table 12.

#### 4.5.2 H-TICK and H-cTICK

The two clocks developed on the human data followed a very similar optimization process with respect to those developed for *N. Furzeri*. In these cases, however, the amount of samples available was substantially larger, allowing us to explore larger models, with an ***F****_i_* with a higher number of features. Also, in the H-TICK and H-cTICK the *selector* layer was turned off, since the input data were the 293 human orthologs of the genes selected by the N-TICK and N-rTICK.

Table 13 reports the explored range of the hyperparameters of the model and their actual value in the H-TICK and H-cTICK developed.

**Table 13:** Parameter space explored for the clocks developed on the human data.

| Parameter | Range | H-TICK | H-cTICK |
| --- | --- | --- | --- |
| Neurons first layer ($\mathbb{F}\mathbb{E}$) | 50 – 200 | 256 | 256 |
| Total number of parameters | 10,000 – 1,000,000 | 200,000 | 200,000 |
| Hidden layers ($\mathbb{F}\mathbb{E}$) | 3 – 8 | 5 | 5 |
| Hiddern layers ($\mathbb{C}$) | 3 – 8 | 3 | 3 |
| Hidden layers ($\mathbb{P}$) | 3 – 8 | 3 | 3 |
| Dropout rate | 0.01 – 0.30 | 0.1 | 0.1 |
| Number of learned features ($\mathbb{F}\mathbb{E}$) | 10 – 200 | 60 | 60 |
| Learning rate ($\mathbb{F}\mathbb{E}$) | $10^{-9}$ – $10^{-4}$ | $1 \cdot 10^{-6}$ | $2.5 \cdot 10^{-6}$ |
| Learning rate ($\mathbb{C}$) | $10^{-9}$ – $10^{-4}$ | $1 \cdot 10^{-7}$ | $1 \cdot 10^{-7}$ |
| Learning rate ($\mathbb{F}\mathbb{E}+\mathbb{P}$) | $10^{-9}$ – $10^{-4}$ | $2.5 \cdot 10^{-6}$ | $5 \cdot 10^{-6}$ |
| $\gamma$ | — | — | — |
| $t$ | — | — | — |
| Batch size | 10 – 1000 | 300 | 300 |

**Table 14:** Repository references for the datasets used.

| Dataset ID | Dataset name | Repositories |
| --- | --- | --- |
| Dataset 1 | Multi-tissue dataset | GSE52462<br>GSE66712<br>GSE72584<br>GSE104321<br>GSE103816<br>GSE117806<br>GSE103819<br>GSE207748 |
| Dataset 2 | Longitudinal dataset | GSE66712<br>GSE150318 |
| Dataset 3 | GRZ dataset | GSE207748<br>GSE103137 |
| Dataset 4 | Rotenone's dataset | GSE103819<br>GSE103809<br>GSE103805<br>GSE103139<br>GSE66712 |
| Dataset 5 | Wild fishes dataset | GSE183039<br>GSE183037 |

#### 4.5.3 Timing and computational resources

For each of the clocks developed, the training process was interrupted after 1,000 epochs and the whole optimization phase took roughly 2 weeks on an Nvidia 1080 TI GPU.

### 4.6 GO annotation heterogeneity analysis

To perform our permutation test to verify if the genes selected by our N-TICK and N-rTICK were more heterogeneous in terms of their functional role, we used the acyclic GO graphs of the ”Biological process”, ”Molecular function” and ”Cellular component” domains in humans from http://geneontology.org/docs/download-ontology/. The edges of these graphs can be of 5 types: ”is a”, ”part of”, ”regulates”, ”negatively regulates” and ”positively regulates”. We took into consideration only the first two types of edge because they are the only ones that indicate a parental relationship. To effectively measure the heterogeneity of the gene lists we needed to avoid counting multiple times GO terms with a very similar meaning such as “regulation of programmed cell death” and “negative regulation of cell death”. Thus, we decided to cut each GO tree at layer 3, which contains broad GO categories that are substantially different from each other. Any GO term associated to our genes that occupies a layer deeper than 3 is converted to its ancestor at layer 3, while all the GO terms occupying a layer higher than 3 are not modified. The total number of unique GO terms obtained in this way indicates how many ”truly” different functions are represented in the original list of genes.

The permutation tests shown in Fig.s 6 and 8 were performed by calculating this number also for the randomly-generated lists of genes, and comparing it to that of the genes selected by our N-TICK and N-rTICK.

## 5 Data and code availablity statement

The data of *N. furzeri* fish used in this study have been generated within our facility at the Leibniz Institute on Aging – Fritz Lipmann Institute and most of them were progressively published in different GEO repositories. A dataset containing newly acquired samples has been deposited in NCBI’s Gene Expression Omnibus [76] and will be accessible through GEO Series accession number GSE207748 once the paper will be accepted for publication in a peer-reviewed journal.

## 6 Acknowledgement

The Core Facility DNA sequencing of the FLI is gratefully acknowledged for their technological support in library preparation and sequencing. Furthermore, the excellent technical assistance of Ivonne Heinze and Ivonne Goerlich is gratefully acknowledged. We acknowledge the support of the Fish Facility of the FLI.

## Appendix A Supplementary information

### A.1 Limitations of the filtering layer

Our experience has shwon that the filtering layer as implemented has some limitations.

First, its selected genes depend highly on the random seed as well as the number of epochs selected for training. This is somewhat expected, as it is very likely that the information on aging is spread in a redundant way through the transcriptome; however, it remains an issue as it makes the findings less reproducible. Second, the number of genes tend to go down as the number of epochs increases, and one needs to be very careful in setting a high enough threshold for the L1 norm, to avoid remaining with 0 genes. As already explained in the text, this is an artifact of the trick used for gradient calculation. Third, the threshold of the L1 norm can be used to influence the number of genes selected (a higher threshold will result in more genes selected, as it allows for a larger sum of weights). However, it is imprecise and again difficult to implement reliably, given that the number of genes also depend on the length of the training process.

### A.2 Effectiveness of the adversarial learning framework

In order to show that the proposed adversarial learning approach indeed eliminates the effect of confounders from the features extracted by FE, we report the correlation between the prediction of C and the true value for each confounder. Fig. A1a shows the correlation when the adversarial learning framework is switched on, while A1b shows it while it is switched off. It can be noted that the influence of the confounders is greatly decreased when the adversarial learning is enabled. We reported the case of the N-TICK, but similar trends can be observed for the other predictors too.

### A.3 Error analysis of N-TICK, N-rTICK, H-TICK and H-cTICK across different stratifiers

To assess the generalizability of our clocks, we report a set of box plots showing the prediction errors for different stratifiers of the datasets. Fig. A2 and A3 show the residuals of the N-TICK predictions on dataset 1 and 2, respectively. Fig. A4 and A5, instead, show the N-rTICK errors on dataset 1 and 2, respectively. Using the same data visualization approach, Fig. A6 shows the residuals of the H-TICK, while Fig. A7 reports the percentage of misclassified subjects for different stratifiers.

As it can be noted, there are no significant differences among the various subcategories of data in any of the box plots illustrated. The only exception is represented by the distribution of the N-rTICK errors across the batches of dataset 1, but we assessed that the batches with higher errors where those composed by samples acquired at a higher age and, as explained in 1.7, the N-TICK errors increase with age.

Finally, for the sake of completeness, we report the confusion matrices of the H-cTICK computed on the training and validation set, given that in the main text we only reported those computed on the test and hold-out sets. As discussed for the other two confusion matrices there are no particular patterns observable in the error distribution of the H-cTICK.

### A.4 Analysis of the N-rTICK sensitivity to genetic, environmental and treatment based lifespan modulating factors

Fig. A9 summarizes the analyses illustrated in Fig. 2 and 3 performed, instead, using the N-rTICK. In all these analyses we can see that the N-rTICK is still sensitive to genetic, environmental and treatment based lifespan modulating factors. In fact, the age of the GRZ and wild samples are overestimated and diverse effect of Rotenone at 9 weeks and 27 weeks are still detectable. However, by making a direct comparison with the results obtained using the N-TICK (i.e. Fig. 2 and 3), important differences are evident:

- the distribution of the errors is more spread;
- in Fig. A9c the slope of the red line is negative, meaning that according to the N-rTICK predictions the biological age of the wild individuals is decreasing along time;
- the rejuvenating effect of Rotenone on the oldest fish is barely recognizable, while the pro-aging effect on the youngest population is detectable but less pronounced. Thus, the N-rTICK recognizes the effect of one type of lifespan manipulation but not its opposite.

While the error increase was somewhat expected considering that the N-rTICK uses less parameters, the last two points indicate that the N-rTICK predictions can not be considered reliable even within a certain degree of uncertainty.

### A.5 Data visualization

Many figures in this work are scatter plots displaying the predicted age on the y axis and the chronological age on the x one (Fig. 1b, 2b, 3a, 4b, A9a, A9c). However, visualizing each point can be difficult, especially when they overlap as in the case of the N-TICK predictions that are very precise. Thus, in Fig. A10 and A11 we report the same figures using a violin plot representation. Fig. 10b, instead, shows the violin plots of the training, validation, test and hold-out sets H-TICK predictions. To ease its interpretation, we also report in Fig. A12 a violin plot representing all these sets together.

Finally, we report in Fig. A13 two Venn diagrams showing the intersections between the genes analyzed, those selected by our N-TICK and N-rTICK and those that have a human ortholog.

---

**Fig. A1:**
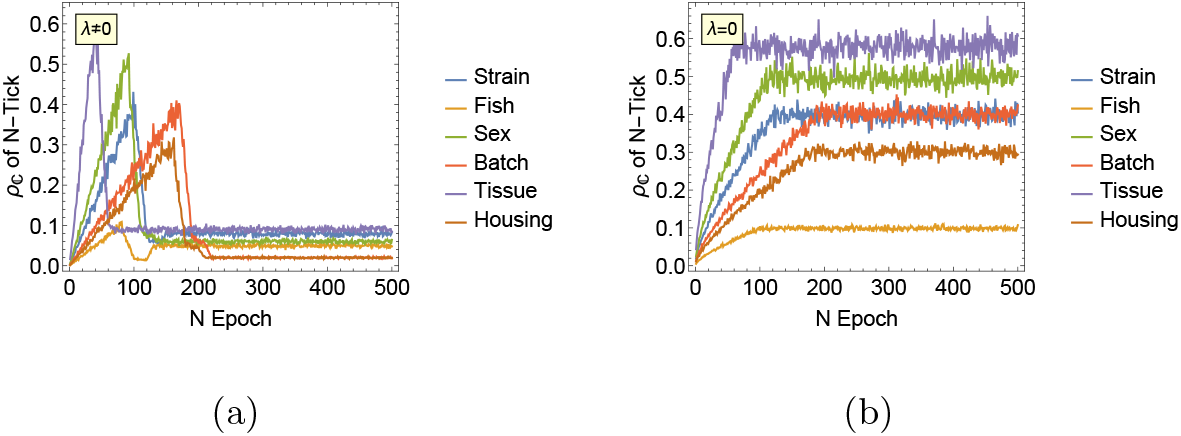
Correlation between the predicted and true value of different confounders across different training epochs, when the adversarial learning framework is switched on (a) and off (b).

**Fig. A2:**
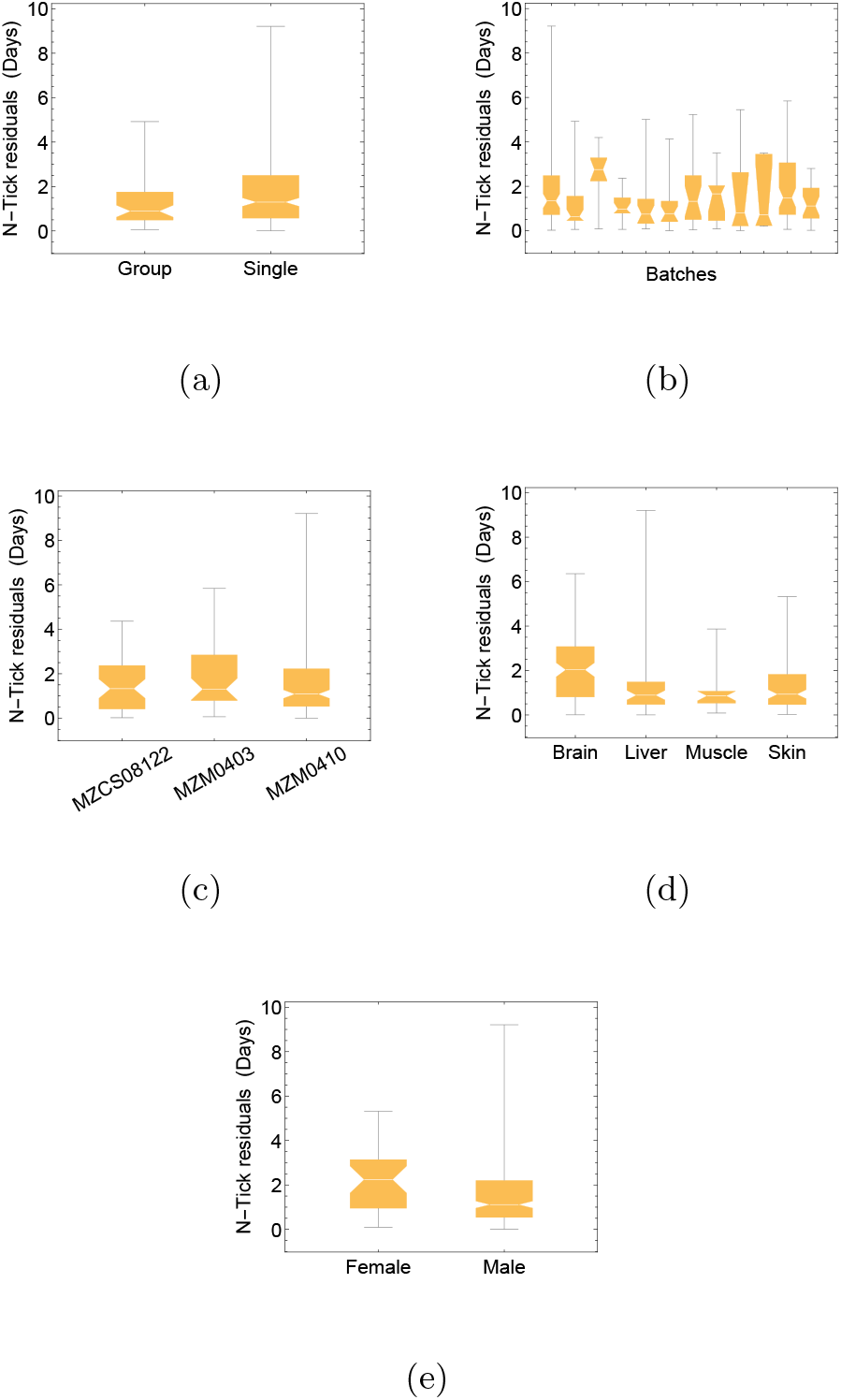
Residuals of the N-TICK predictions on the multi-tissue dataset (dataset 1) for different stratifiers of the dataset: strain (a), tissue (b), sex (c), housing (d) and batch (e).

**Fig. A3:**
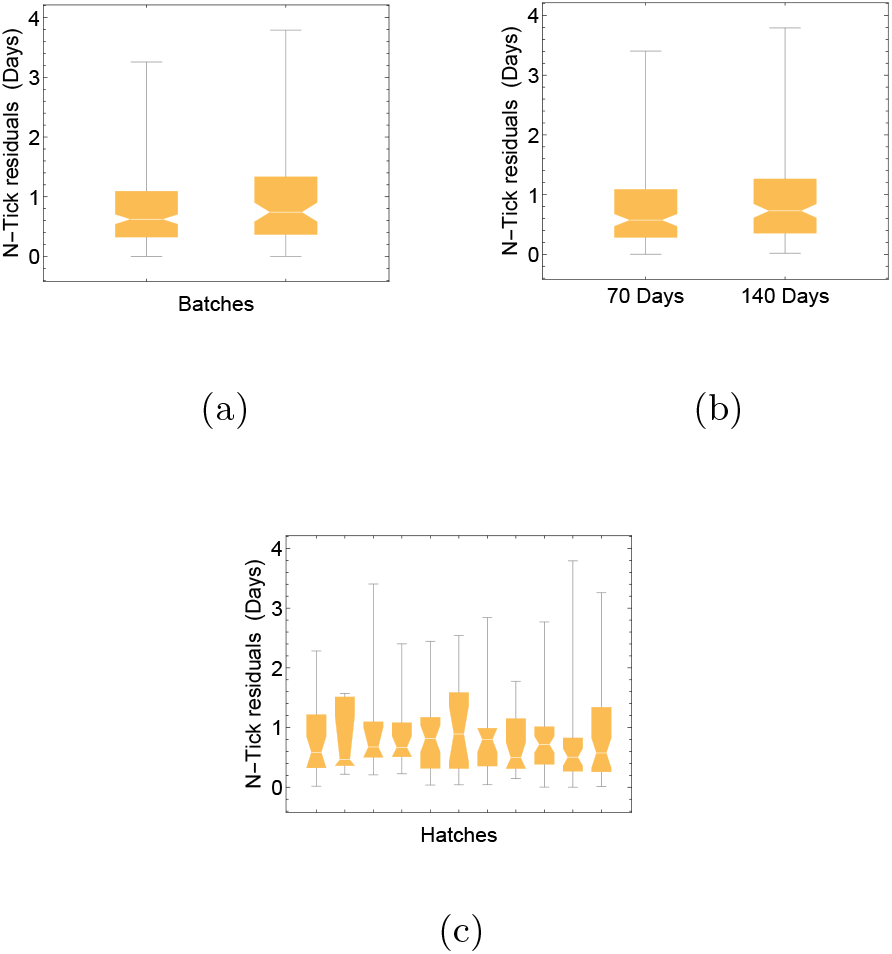
Residuals of the N-TICK predictions on the longitudinal dataset (dataset 2) for different stratifiers of the dataset: batch (a), age (b) and hatches (c).

**Fig. A4:**
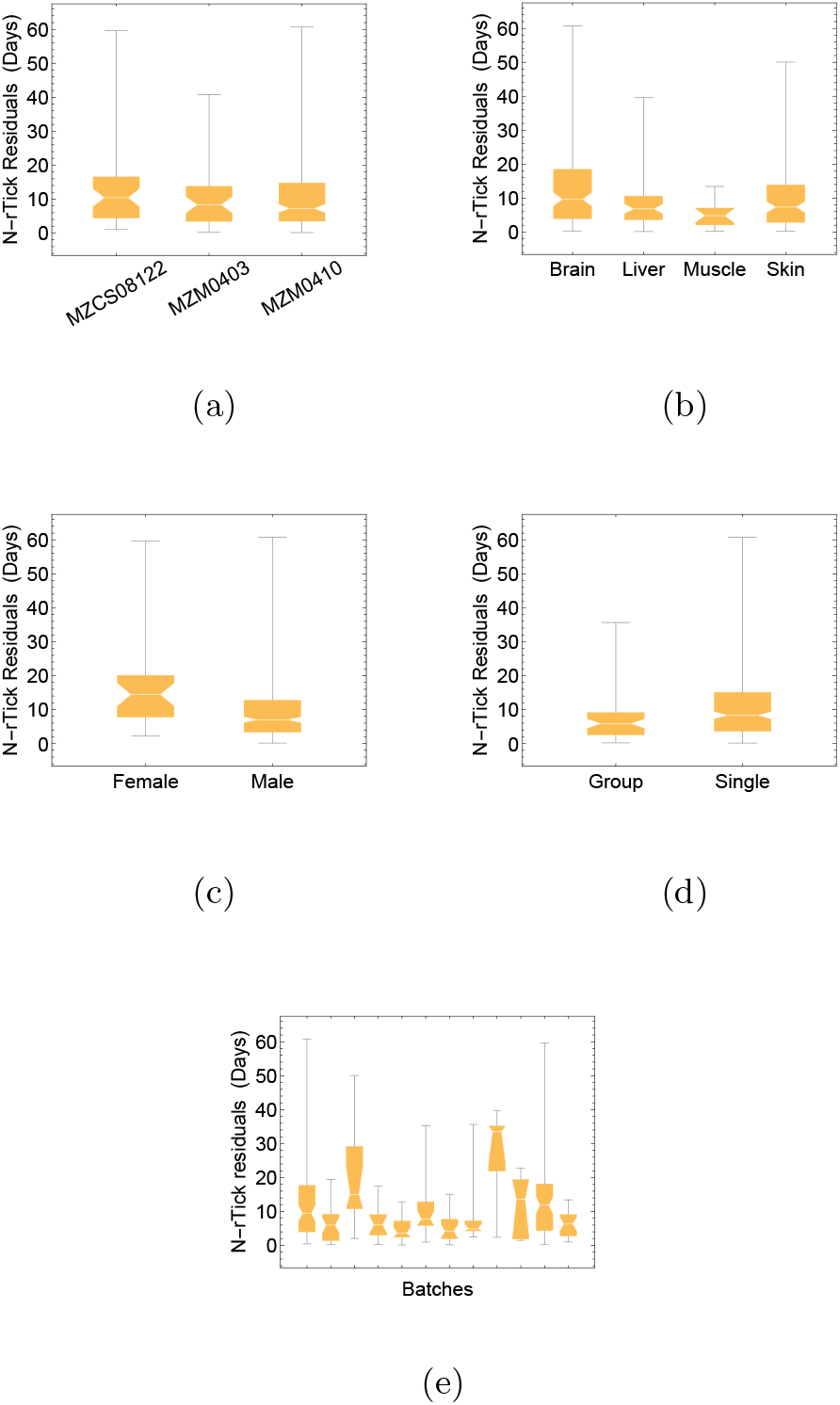
Residuals of the N-rTICK predictions on the multi-tissue dataset (dataset 1) for different stratifiers of the dataset: strain (a), tissue (b), sex (c), housing (d) and batch (e).

**Fig. A5:**
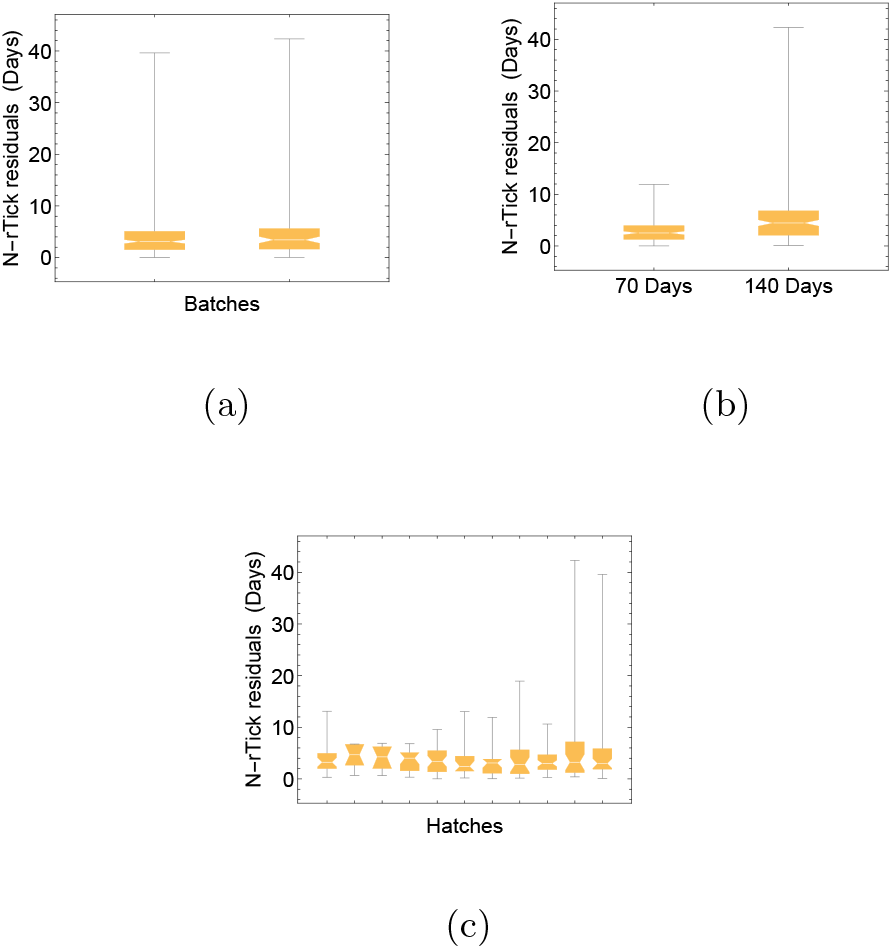
Residuals of the N-rTICK predictions on the longitudinal dataset (dataset 2) for different stratifiers of the dataset: batch (a), age (b) and hatches (c).

**Fig. A6:**
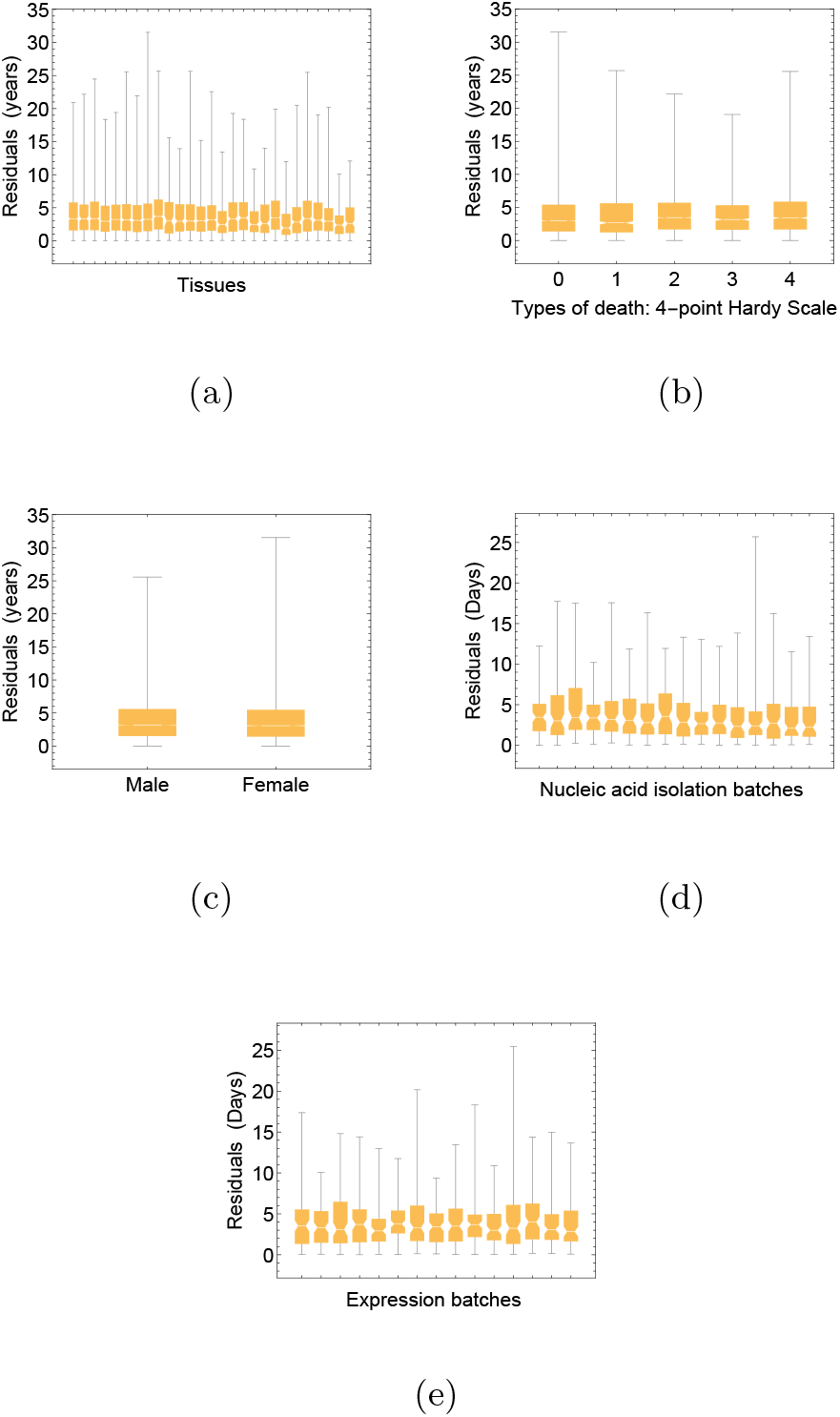
Residuals of the H-TICK predictions divided for different stratifiers of the dataset: tissue (a), type of death (b), sex (c), nucleic acid isolation batch (d) and expression batch (e).

**Fig. A7:**
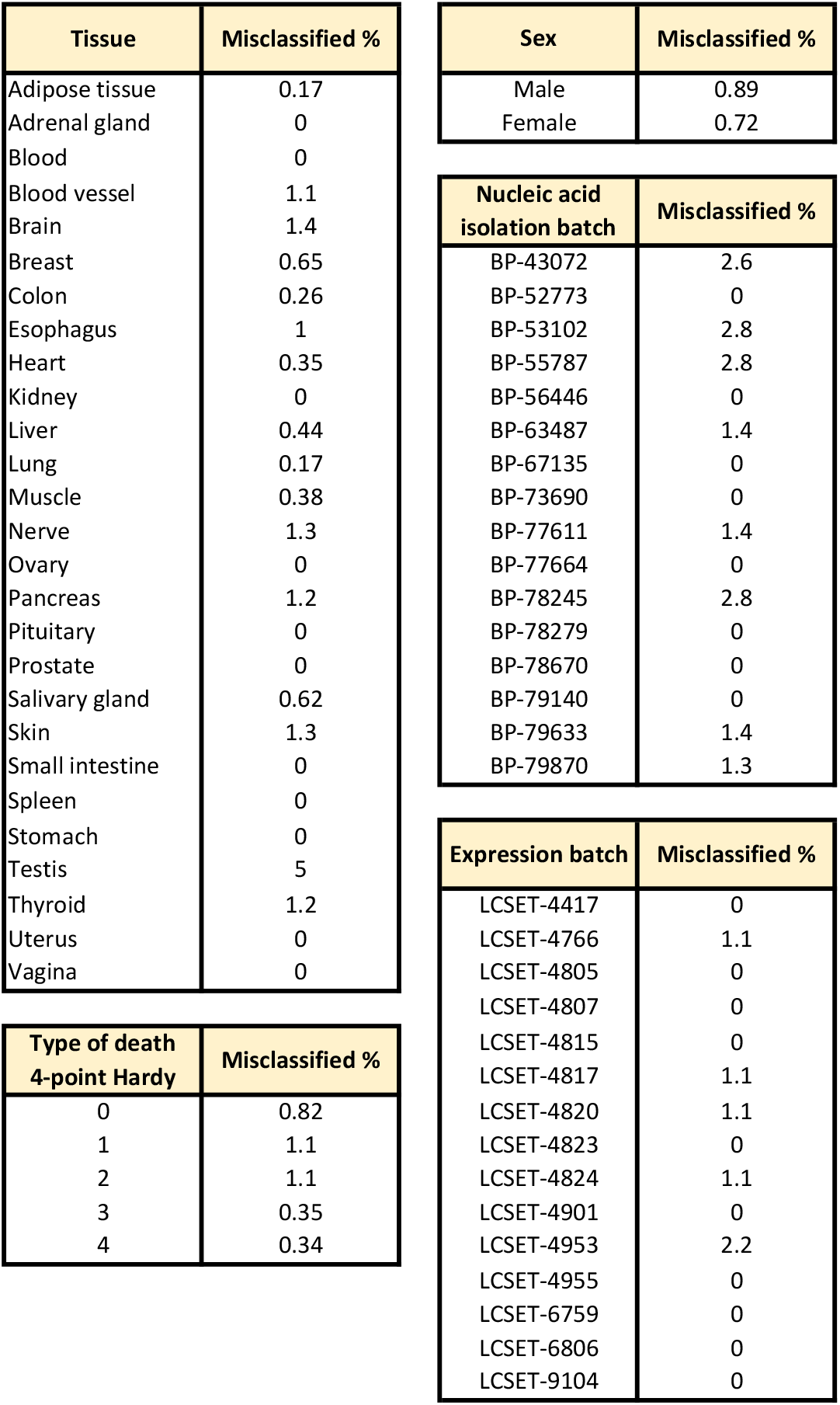
Percentage of H-cTICK mislassified samples divided for different stratifiers: tissue (a), type of death (b), sex (c), nucleic acid isolation batch (d) and expression batch (e).

**Fig. A8:**
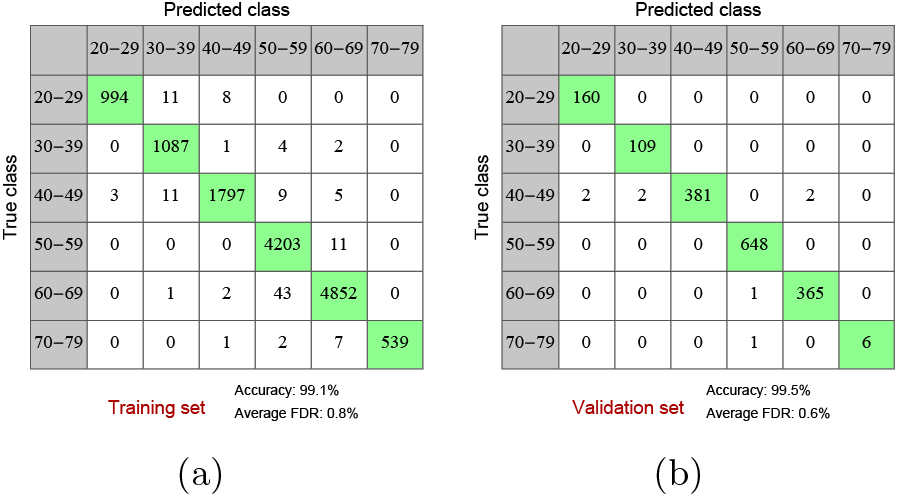
Confusion matrices of the H-cTICK on the training and validation set.

**Fig. A9:**
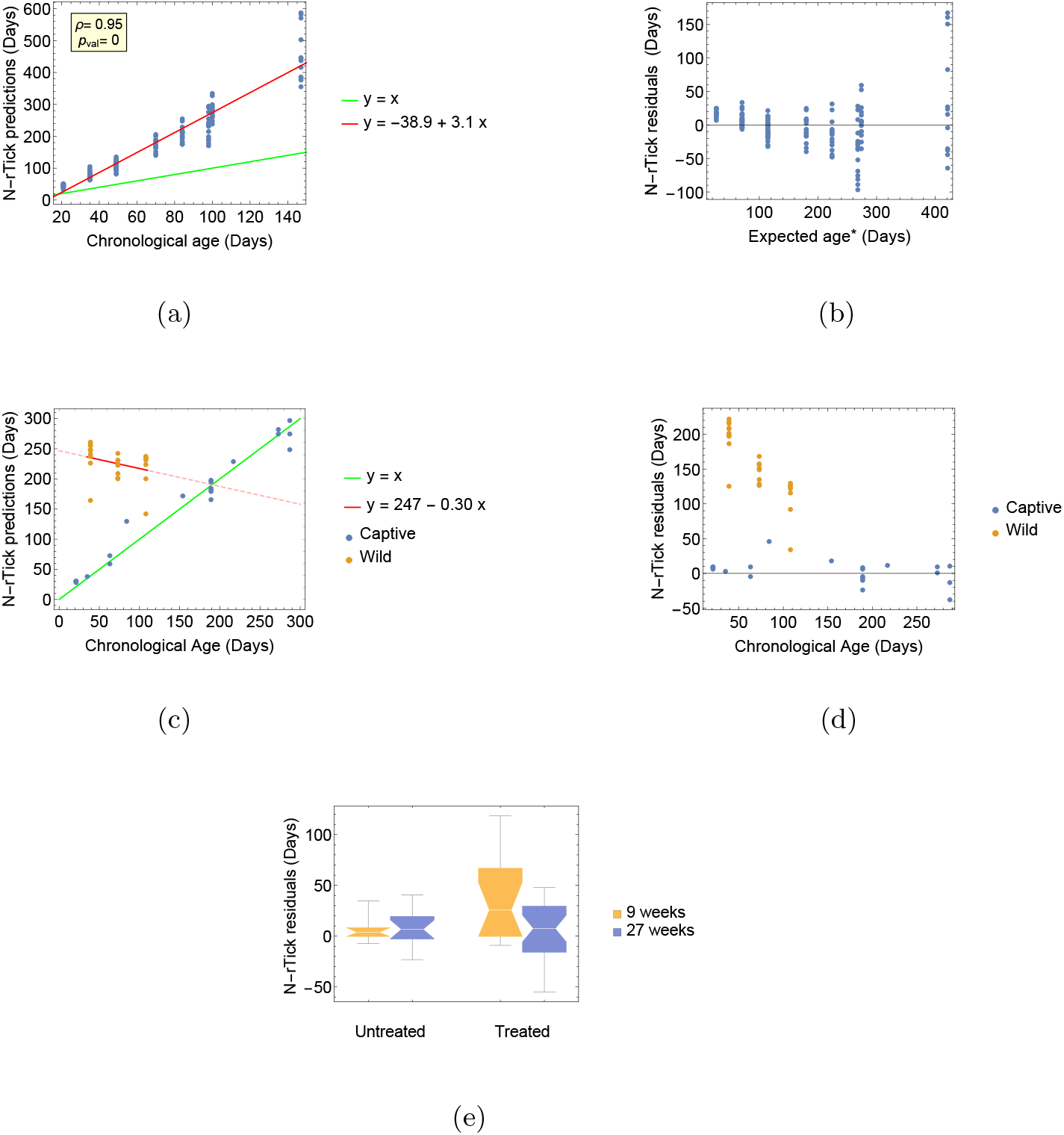
Other results obtained with the N-rTICK. Fig. (a) shows the N-rTICK predictions on the GRZ dataset (dataset 3) and Fig. (b) shows its residuals with respect to the expected age, i.e. the one fit with the red line in Fig. (a). Fig. (c) shows the N-rTICK predictions on brain samples of wild *N. furzeri* fishes (orange dots) and of captive ones (blue dots) and Fig. (d) shows their residuals. Fig. (e) shows the residuals of the N-rTICK predictions on samples treated and untreated with Rotenone.

**Fig. A10:**
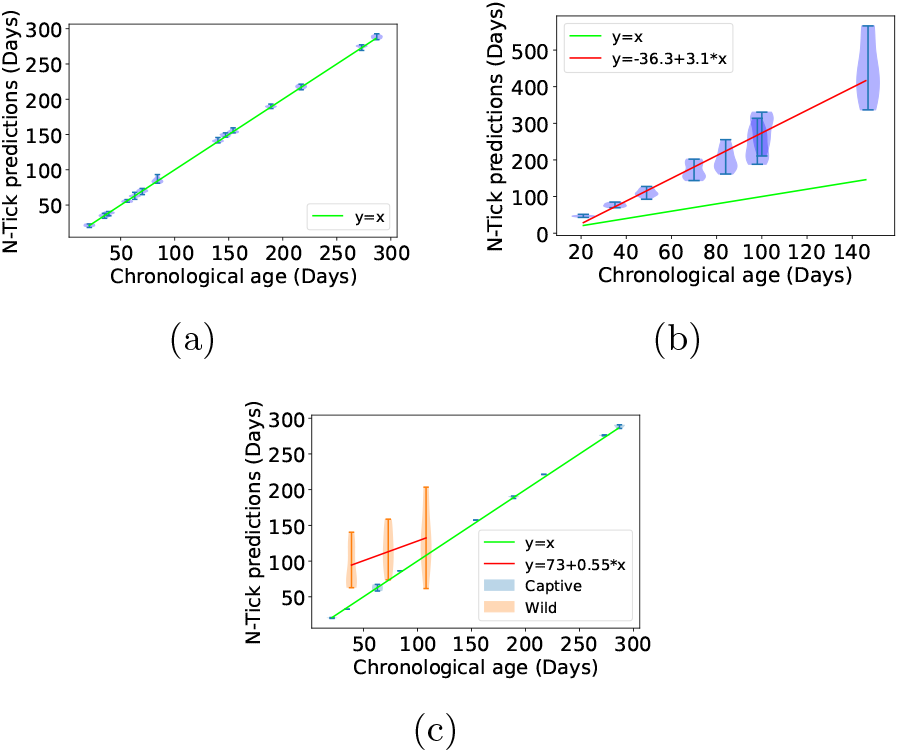
Violin plots of the N-TICK performance. Fig. (a), (b) and (c) show the N-TICK age predictions vs the true age of the samples on datasets 1-2, 3 and 5, respectively.

**Fig. A11:**
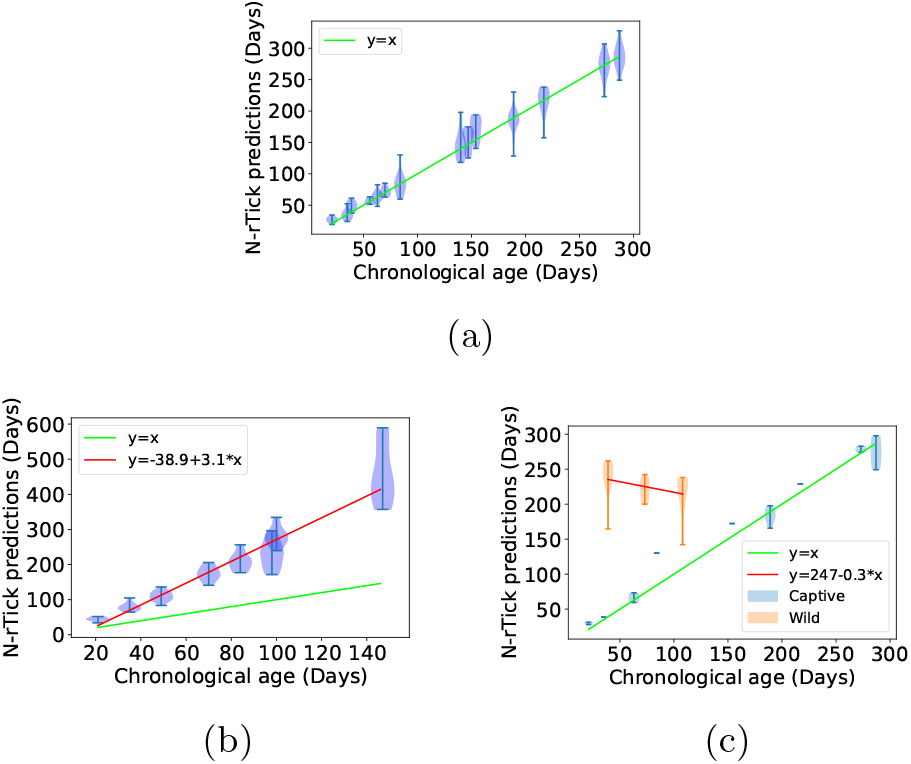
Violin plots of the N-rTICK predictions on dataset 1 (Fig. a) on the GRZ dataset (Fig. b) and on brain samples of wild and captive *N. furzeri* fish (Fig. c).

**Fig. A12:**
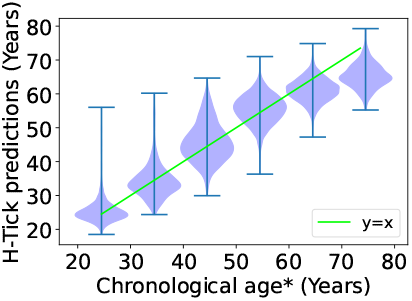
Violin plot of the H-TICK predictions.

**Fig. A13:**
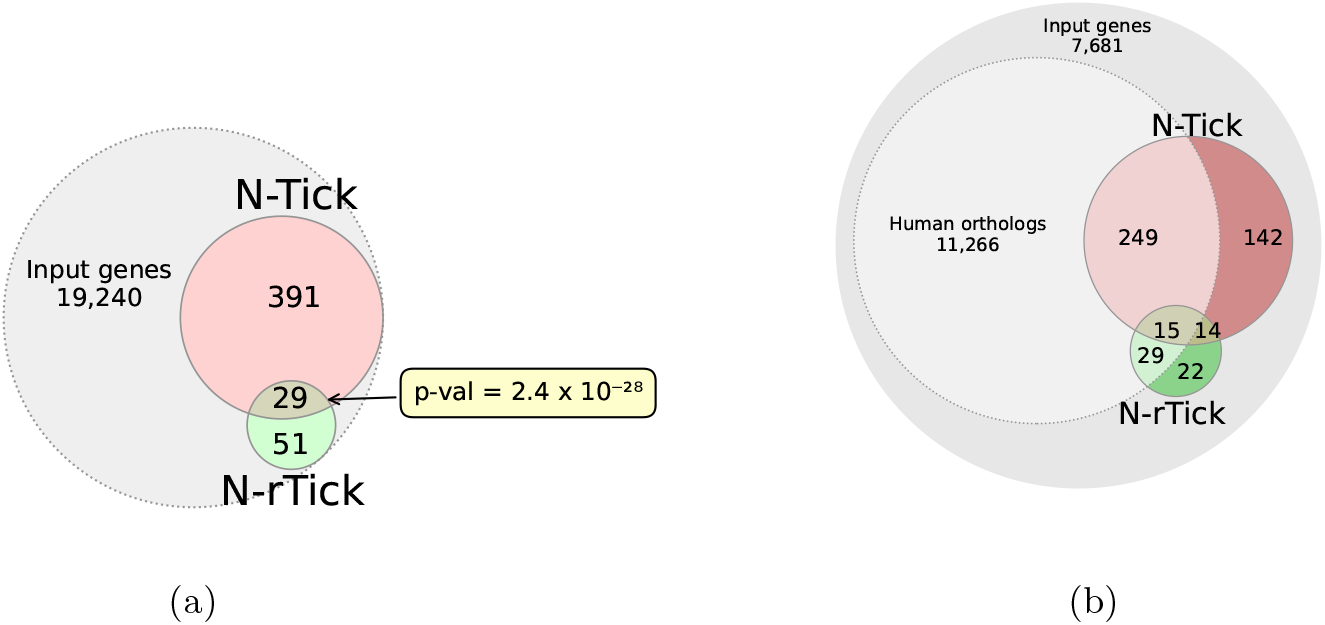
Venn diagrams of the gene list found in this study. Fig. (a) shows the intersections between the genes selected by N-TICK, the ones selected by N-rTICK and the input gene list. Fig. (b), in addition to (a), also shows the human orthologs set.

## Footnotes

1 Errors on categorical and on continuous confounders are unavoidably incomparable. Despite this, we empirically noticed that with these choice of error metrics, the overall loss is able to correct both kinds of confounders.

